# Complementation of a *Setaria* Rubisco activase mutant with *Agave* Rubisco activase restores growth and photosynthesis

**DOI:** 10.64898/2026.08.28.747894

**Authors:** Amber M. Hotto, Sarah Gartner, Kathryn Eshenour, David B. Stern

## Abstract

Photosynthetic carbon assimilation is sensitive to heat stress, with a key component being the thermal sensitivity of Rubisco activase (RCA). One approach to increasing heat tolerance of photosynthesis is to increase the thermotolerance of RCA. Here, we have used a transgenic approach to express RCAβ from *Agave tequilana* (AtRCA) in *Setaria viridis*. A second line was created where the endogenous SvRCAβ was substituted for AtRCAβ through complementation of a null mutant, ΔrcaB (AtRCA[bKO]). *In vitro* assays showed that Agave RCA readily activates *Setaria* Rubisco, and that its thermostability is higher than that of *Setaria* RCA. AtRCA[bKO] plants exhibited higher CO_2_ assimilation at 25°C compared to WT and AtRCA plants, although Rubisco content and activation did not change. After six hours at 44°C, AtRCA[bKO] plants retained higher CO_2_ assimilation rates than AtRCA, however, after 24 hours CO_2_ assimilation declined to similar levels in all lines, suggesting that RCA may not limit carbon assimilation at the conditions tested. Taken together, our results show that substitution of RCA from a CAM plant into a C_4_ model allows normal growth and supports slightly increased carbon assimilation under control and short-term heat conditions.

**Highlight:** Genetic crosses were used to exchange the native Rubisco activase (RCA) in *Setaria viridis* to one from *Agave tequilana*, a desert plant with a highly thermostable RCA. Agave RCA supported slightly increased photosynthesis at ambient temperatures but not under heat stress.

## Introduction

Ribulose 1,5-bisphosphate carboxylase/oxygenase (Rubisco) is a key catalytic enzyme required for the incorporation of atmospheric CO_2_ into sugars required for plant growth. Rubisco activity can be limited by its propensity for binding inhibitory sugar-phosphates at its active site, including its substrate ribulose-1,5-bisphosphate (RuBP). Rubisco activase (RCA) evolved to maintain Rubisco activity by removing these inhibitors and is nearly ubiquitous in autotrophic organisms. RCA is a member of the AAA+ ATPase family that uses ATP hydrolysis to generate structural changes in Rubisco that facilitate inhibitor diffusion from the active site (Waheeda *et al*., 2023). Transgenic plants depleted for, or mostly lacking, RCA have been examined in several species including Arabidopsis, tobacco, rice, *Flaveria* and *Setaria*, and in all cases this creates a requirement of elevated CO_2_ for survival when RCA becomes sufficiently depleted (Somerville *et al*., 1982; Mate *et al*., 1993; von Caemmerer *et al*., 2005; Hotto *et al*., 2025).

RCA is an important factor in the heat sensitivity of photosynthesis (Sharkey *et al*., 2001), because of the heat lability of its ATPase activity, particularly in C_3_ species (Salvucci *et al*., 2001; Demirevska-Kepova *et al*., 2005; Barta *et al*., 2010; Masumoto *et al*., 2012). 2021). Variation in the thermostability of RCA has also been used as a tool to temper the heat sensitivity of photosynthesis in several species. In Arabidopsis, *in vitro* mutagenesis was used to screen for more heat-tolerant activases, which conferred increased thermotolerance of photosynthesis in transgenic plants when they were substituted for the endogenous RCA (Kurek *et al*., 2007). In wheat, rice and Arabidopsis, genetic crosses or transgenic approaches were used to add new forms of RCA with greater thermotolerance, again partially mitigating heat stress of photosynthesis (Scafaro *et al*., 2018, 2019; Wijewardene *et al*., 2020), including the overexpression of maize RCA in rice (Qu *et al*., Such results have led to suggestions that RCA should be a target for crop improvement, particularly in a climatic era where heat stress is expected to be more frequent and severe (Ogbaga *et al*., 2018; Wijewardene *et al*., 2021; Gjindali *et al*., 2025).

While complementation of a null mutant was used in some Arabidopsis studies to exchange RCAs, in most of the studies cited above divergent RCAs were added to WT plants resulting in the co-expression of multiple RCA isomers that would be expected to form heterooligomers with intermediate properties (Zhang *et al*., 2001; Shivhare and Mueller-Cajar, 2017; Shivhare *et al*., 2019). In a recent study, we reported the generation of gene edited mutants for the two forms of RCA in *Setaria viridis*, RCAα and RCAβ (Hotto *et al*., 2025). In *Setaria*, RCAβ is the only RCA form present at 25°C, therefore, the mutant, which we termed bKO, is a null RCA mutant at ambient temperatures and requires elevated CO_2_ for survival. RCAα is induced at elevated temperatures and is thought to stabilize RCA activity under these conditions. Since only RCAβ is expressed in the absence of heat stress, we have complemented the bKO line through genetic crosses that introduce a transgene expressing Agave RCAβ. Agave RCAβ was chosen for its *in vitro* thermostability of ATPase activity (Shivhare and Mueller-Cajar, 2017), thus potentially allowing us to modify the heat response of carbon assimilation in *Setaria viridis.* While plants expressing Agave RCAβ in place of *Setaria* RCAβ did not overcome heat sensitivity of photosynthesis after 24 hr, the Agave enzyme readily activated *Setaria* Rubisco and supported higher assimilation rates at ambient temperature. These results illustrate a rigorous approach to testing the importance of RCA variants whose *in vitro* properties suggest a possible functional advantage *in planta*.

## Materials and methods

### Purification and analysis of recombinant RCA

Cloning of *Setaria* RCAβ into pHUE was previously described (Hotto et al., 2025). For this study, the pHUE-AtRCA plasmid expressing amino acids 58-435 from *Agave tequilana* RCAβ was a kind gift from Oliver Mueller-Cajar (Shivhare and Mueller-Cajar, 2017). The *Zea mays* RCAβ amino acids 52-434 were cloned into pHUE as previously described (Catanzariti *et al*., 2004) via SacII and HindIII restriction digest of the PCR product obtained from maize cDNA with primers listed in Table S1. Purification of recombinant RCA was performed as described (Hotto *et al*., 2025), with at least two individual preparations of each RCA (biological replications) to account for variability between preparations.

### ATPase and rubisco activation assays

ATPase assays of recombinant RCAs were performed as described in Hotto et al. (2025), with the following modification: RCA samples were supplemented with 0.2 mM ATP during pre-incubation as addition of ATP is known to increase the thermostability of RCA (Scafaro *et al*., 2019). This was also the case when we compared the thermostability of SvRCA and AtRCA with and without the addition of ATP in an ATPase assay (Fig. S1). Following a 10 min pre-incubation with the designated RCA isoform and ATP at the indicated temperature, the absorbance at 340 nm was monitored for 10 min with data recorded every 10 s. Thermal shift assays were conducted as described (Scafaro *et al*., 2019), with Tm analysis performed on DSF world (https://bio.tools/dsfworld) as described (Wu *et al*., 2020).

Rubisco activation assays were performed as described (Oh et al., 2024) with the following modifications: reactions were initiated upon addition of either ECM (fully activated) or ER (RuBP-inhibited) Rubisco; ER was prepared by adding the same ratio of extract used to make ECM (70% of total volume); and absorbance was recorded for 6 min. ECM and ER were generated from wild-type *Setaria viridis* Rubisco extracts quantified as described previously (Eshenour et al., 2024). To determine RCA’s ability to activate ER, 5 µM RCA were added to the assay. Results of the assays measured the ability of recombinant RCA to reactivate extracted Rubisco following its *in vitro* inhibition by incubation with RuBP.

### Generation and identification of Agave tequilana (At) RCA expressing Setaria plants

To obtain *Setaria* plants expressing chloroplast-localized *Agave tequilana* (At) RCAβ, subsequently referred to as ‘RCA’, the AtRCA gene was amplified from the pHUE-AtRCA vector (described above) using gene-specific primers (Table S1). The maize RCA transit peptide, amplified from maize cDNA, was added to the 5’ end of the AtRCA gene via overlap extension PCR. The Zm_TP_-AtRCA PCR product was directionally cloned into pENTER/D-TOPO. The final construct was cloned into the pANIC10A vector (Mann *et al*., 2012) using Gateway technology and verified by sequencing and restriction digest. The pANIC10A-AtRCA plasmid was transformed into *Setaria* ME024 at The Center for Biotechnology Research at the Boyce Thompson Institute, with positive transformants identified by growth on media supplemented with hygromycin. The T3 generation was identified as homozygous when 100% of the progeny contained the AtRCA transgene by PCR with the primers listed in Table S1.

Homozygous AtRCA lines lacking endogenous SvRCA were generated via genetic crosses to ΔrcaB (bKO) plants according to Jiang et al. (2013). For these crosses, bKO plants were emasculated and pollinated with AtRCA expressing lines. Seeds were germinated and grown at 9,000 ppm CO_2_ for 2-3 weeks, after which the plants were moved to ambient air (400 ppm CO_2_). Products of successful crosses were able to survive in ambient air, while seedlings derived from uncrossed ΔrcaB could not. PCR was used to verify the presence of the AtRCA transgene in surviving plants, and the presence of the inactivating gene edit in the native *Setaria RcaB* gene was checked via PCR, sequencing and comparison to the WT *RcaB* gene using ICE analysis (Hotto et al., 2025; Figs. S2A and S2B). The event predominantly analyzed here, AtRCA [bKO], contained 100% gene editing determined by sequencing along with the presence of the Agave transgene. A second AtRCA transformation event was also crossed to bKO and, while all plants contained the Agave transgene (Fig. S3A), approximately 30% of the sequenced portion of the *Setaria RcaB* gene was a wild-type sequence based on ICE analysis and sequencing of individual *RcaB* clones (Fig. S3B). While we are uncertain of the source of these WT sequences, their proportion is stable between generations, suggesting that a single locus may contain both the edited WT gene and a fragment of the WT gene that appears to have been present in that second AtRCA event prior to the genetic cross.

For PCR, DNA was extracted from leaf tissue by grinding tissue into a powder, suspension in lysis buffer (250 mM NaCl, 25 mM EDTA, 0.5% SDS, 200 mM Tris pH 8.0), incubation at 65°C for 15 min, and centrifugation for 5 min at 16,000 x *g*. The supernatant was removed and added to an equal volume of isopropanol, vortexed, and left at room temperature for one hr. Samples were centrifuged at 16,000 x *g* for 15 min, the pellet was washed twice with isopropanol and then resuspended in 1xTE buffer. PCR was completed using GoTaq polymerase (Promega) according to the manufacturers’ instructions.

### Plant growth conditions

Seeds aged at least four weeks postharvest were sterilized with 10% bleach and 0.1% Tween-20 for 5 min with agitation, washed three times with sterile water, then sown on MS agar. Seedlings were transplanted to soil (BK25 supplemented with lime and unimix) in 6 cm pots 5-7 days after sowing (DAS), then into 15 cm pots 21 DAS, with fertilization at least three times weekly. Plants were grown in growth chambers at 25°C with a 16:8 h light/dark cycle, 50% humidity and 300 µmol m^−2^ s ^−1^ light intensity. Control measurements were taken at 25-29 DAS. For heat treatment, plants were kept at 44°C for up to 24 hours. Results shown were collected from at least three batches of plants containing all three genotypes, grown at separate times under the same conditions. Plant phenotypes were taken 14, 21, 28 and 35 DAS under control conditions. Plant height, tiller count and dry weight were measured as described (Hotto et al., 2025).

### Protein isolation and analysis

Proteins were isolated on an equal leaf area basis. Tissue was harvested from the middle portion of the youngest fully expanded leaf, frozen in liquid N_2_ and stored at -80°C. For analysis, tissue samples were ground to a powder in liquid N_2_ and suspended in equal volumes of 2x Laemmli buffer with 0.015% DTT (w/v). The samples were vortexed, heated at 65°C for 15 min, then centrifuged at 16,000 x *g* for 30 sec. The supernatant was loaded in 8% SDS-polyacrylamide gels, then transferred to methanol-activated polyvinylidene fluoride membranes (Cytvia, Marlborough, MA, USA). Membranes were blocked in a Tris-buffered saline/0.1% Tween-20 (TBS-T) and 5% (w/v) milk solution overnight at 4°C with shaking, and incubated with primary antibodies in TBS-T and 5% (w/v) milk at room temperature for 1 h. Primary antibodies from Agrisera (Vännäs, Sweden; www.agrisera.com) were anti-RCA (1:10,000), anti-AtpB (1:10,000), anti-PPDK (1:5,000), anti-LS (1:20,000) and anti-CytF (1:5,000). Membranes were rinsed three times in TBS-T for 10 min, then incubated in secondary antibody (1:10,000 goat-antirabbit IR dye 800 CW; LI-COR Biosciences) in TBS-T and 5% (w/v) milk for 1 h at room temperature, rinsed three times in TBS-T for 10 min and once in TBS for 10 min, then imaged using the LI-COR Odyssey Infrared Imaging System. Membranes were stained with 0.01% Coomassie Blue R-250, destained with 7% acetic acid/40% methanol solution, and imaged.

### Mesophyll and bundle sheath cell extractions

Mesophyll and bundle sheath (BS) separation used up to 5 g leaf tissue from mature plants using only the middle section of fully expanded leaves, discarding both the leaf tip and the tissue close to the collar. Mesophyll protoplasts were isolated as described previously (Hotto et al., 2021) with the final pellet resuspended in 500 µL of wash buffer and stored at -80°C until use. BS samples were prepared largely as described (Markelz *et al*., 2003). Briefly, leaf tissue was cut transversely into small strips then pulsed in a blender with dull blades and BS buffer 1 (0.33 M sorbitol, 0.3 M NaCl, 0.01 M EDTA, 0.01 M DTT, 0.2 M Tris, pH 9.0) at low speed for 3 x 10 sec. The mix was filtered through a 60 µm nylon mesh, and the remaining leaf debris was pulverized at high speed in a blender for 1 min in bundle sheath buffer 2 (0.35 M sorbitol, 5 mM EDTA, 0.1% β-mercaptoethanol, 0.05 M Tris, pH 8.0). This was repeated three times, and the final tissue was dried slightly and stored in batches wrapped in foil at -80°C until use.

### Photosynthesis measurements

Gas exchange measurements were made with an LI-6800 Portable Photosynthesis System (LI-COR, Lincoln, NE, USA) with a 2 cm^2^ leaf chamber. Measurements were taken from the youngest fully expanded leaf on the primary tiller. All measurements assessed genotypes in a randomized order between 2 h and 8 h into the light period. To measure CO_2_ assimilation rates (*A)* at saturating light and ambient CO_2_ (*A*_sat_; C_a_ [concentration of CO_2_ at leaf level] of 400 µmol mol^−1^) and assimilation rate at saturating light and maximum CO_2_ (*A*_max_; C_a_ of 1500 µmol mol^−1^), plants were acclimated to ambient CO_2_ (400 µmol mol^−1^), the leaf chamber was set to the desired chamber temperature, and 1500 PPFD light was applied for 15-20 min.

To measure CO_2_ assimilation rates over a range of temperatures, plants were acclimated to ambient CO_2_ (400 µmol mol^−1^) with the leaf chamber set to 34°C, and non-saturating light of 400 PPFD (to avoid photobleaching) was used for 15-20 min. The temperature of the leaf chamber was increased by 2°C every 10 min and *A* values recorded every 10 sec for 110 min. To measure the time course of photosynthetic activation, the leaf chamber was set to 0 PPFD for 30 min, then light was increased to 1500 PPFD for 30 min, with *A* values recorded every 10 sec. The leaf chamber was set to 25°C and ambient CO_2_ for the duration of the experiment. *A* values were normalized using the highest recorded value during the light period (PPFD=1500) for each individual plant and the lowest recorded values in the dark period (PPFD=0).

### Rubisco quantification and activity

Tissue samples for determining Rubisco content and activation were collected from the middle region of the youngest fully expanded leaf in the same region that gas exchange measurements were taken. Samples were collected the day following gas exchange analysis after being equilibrated at saturating light (1500 PPFD), then flash frozen in liquid N_2_ and stored at -80°C until use. Rubisco content was quantified using radiometric methods described previously (Eshenour et al., 2024). Percent Rubisco activation was determined from the ratio of the rate of activity of the initial Rubisco extract (Rubisco without activation or inhibition) to the rate of fully activated Rubisco from the same extract. Rubisco activity was measured using the spectrophotometric assay described above. Sample numbers for each experiment are given in the Figure Legends.

### Statistical analyses

One and two-way ANOVA was performed as described in Eshenour et al. (2024). The Student’s unpaired *t*-test was also used via the GraphPad calculator to test for significant differences with a *p*-value of <0.05.

## Results

### Both Agave and Setaria RCAs are thermostable and can activate Setaria Rubisco in vitro

To establish the comparative properties of the relevant RCAs *in vitro*, we prepared recombinant proteins corresponding to RCAβ from *Setaria viridis* (Sv), *Agave tequilana* (At) and *Zea mays* (Zm; Fig. 1A). Maize was used as a reference because, like *Setaria* and Agave, it is a C_4_ plant, however, its photosynthesis is heat sensitive at 38°C (Crafts-Brandner and Salvucci, 2002) in comparison to *S. viridis* (ca. 42°C; Zhang *et al*., 2025). We first measured the heat sensitivity of ATPase activity for the three enzymes, as shown in Fig. 1B. Compared to the activity at 25°C, SvRCA ATPase activity was stable until 40°C, significantly reduced at 45°C, and nearly absent at temperatures above 45°C. AtRCA ATPase activity was about 25% lower than both Zm and Sv RCAs at 25°C, however, this RCA was the most thermotolerant with activity dropping only at temperatures exceeding 45°C. Given that the initial AtRCA ATPase activity was lower than SvRCA, at 45°C the ATPase activity of SvRCA and AtRCA were analogous. The least thermostable RCA was ZmRCA, which lost 23% of ATPase activity at 40°C. These results confirm that ZmRCA is more thermosensitive than SvRCA and AtRCA, while AtRCA is the most thermostable.

**Fig. 1.**
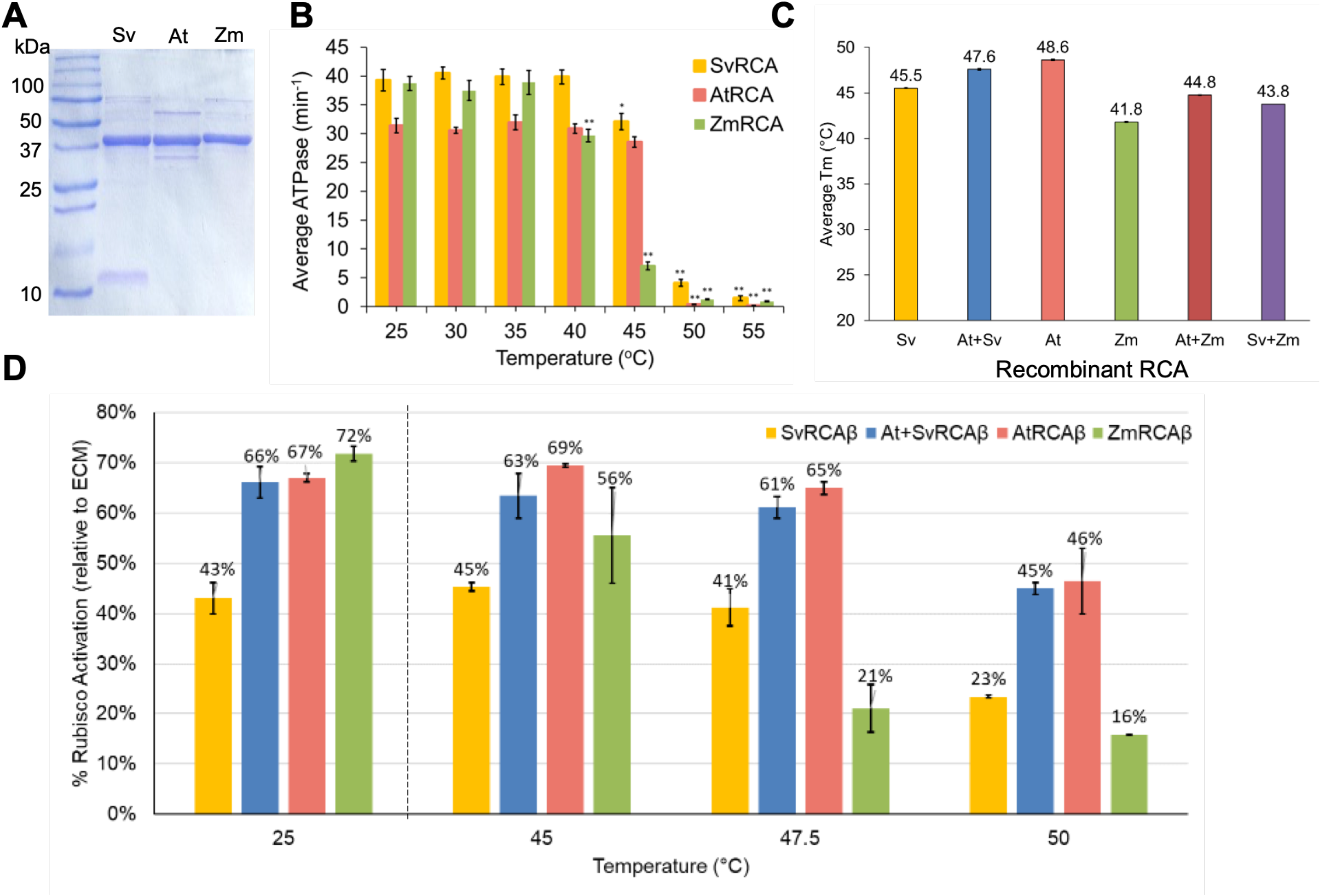
Biochemical characterization of recombinant RCA. (A) Purified recombinant RCAs separated by 13.5% SDS-PAGE and stained with Coomassie brilliant blue. 4 µg of each protein was loaded and the precision plus protein standard was loaded for reference. (B) ATPase activity assays of different RCA isoforms preincubated at the indicated temperature for 10 min, then assayed at 25°C. Significant differences for each RCA isoform compared to the cognizant 25°C activity are indicated (\**p*-value < 0.05; \*\**p*-value < 0.0001) as determined by the Student’s *t*-test. (C) The temperature at which half the structural stability of the indicated RCA isoform, either alone or in a 1:1 molar ratio, is lost (Tm) as determined using a thermal shift assay. (D) Rubisco activation assays by the indicated RCA isoforms individually or in a 1:1 molar ratio preincubated at the indicated temperature, then assayed at 25°C. Data reflect the ability of the RCA isoform to activate fully inhibited Rubisco (ER) relative to fully activated Rubisco (ECM) after 6 min. Data for all assays represent n=2-3 with at least three technical replicates each, and error bars are ± SE.

To determine if the decline in RCA ATPase activity at elevated temperatures was reflected in protein stability, we used a thermal shift assay (TSA) to measure the Tm of individual recombinant proteins as well as 1:1 mixtures that likely form a range of heterooligomers (see below). Figs. 1C and S4 show that the Tm range varied from 41.8°C for ZmRCA to 48.5°C for AtRCA, with SvRCA being intermediate at 45.3°C. These temperatures correlate well with the temperatures at which enzymatic activity decreases (Fig. 1B), suggesting that TSA assays can be reasonably used to predict the temperature at which the loss of ATPase activity will occur. The 1:1 mixtures had values intermediate to their individual components, further suggesting that they formed heterooligomers *in vitro*. Heterooligomers with intermediate properties were previously reported between rice and Agave RCAs (Shivhare and Mueller-Cajar, 2017).

To ensure that the recombinant RCAs could activate *S. viridis* Rubisco, we compared the ability of SvRCA, AtRCA or a 1:1 mixture of to activate inhibited Rubisco (ER) *in vitro* (Fig. 1D). ZmRCA was also included as a thermosensitive reference. All recombinant RCAs assayed at 25°C were able to activate *Setaria* Rubisco. Surprisingly, activation of Rubisco by SvRCA was significantly lower (43%) compared to AtRCA, ZmRCA and At+SvRCA (66-72%). The high degree of *Setaria* Rubisco activation by AtRCA suggests that AtRCA should also be able to substitute for SvRCA *in planta*. To examine the thermostability of the recombinant RCAs in activating Rubisco, each RCA was first incubated at the indicated temperature, then added to fully inhibited Rubisco (ER) and Rubisco activation was monitored. Temperatures were chosen based on the temperature at which RCA ATPase activity was lost and the RCA Tms. For Sv and Zm RCAs, Rubisco activation activity was retained at higher temperatures compared to the temperature at which ATPase activity declined (45°C and 40°C, respectively), with SvRCA activity declining above 47.5°C and ZmRCA activity above 45°C. AtRCA Rubisco activation activity declined significantly at 50°C, however, Rubisco activation was still at 50% vs almost no ATPase activity at the same temperature. This data indicates that ATPase activity is not necessarily reflective of Rubisco activation efficiency. Heterooligomers of Sv and AtRCAs were also examined to mirror what might be happening in transgenic plants expressing both RCAs. In this case, a 1:1 ratio of Sv + At RCAs were significantly better at activating Rubisco than SvRCA alone, both at 25°C and higher temperatures, similar to AtRCA alone. The implications of this result are further considered in the Discussion.

### Expression of AtRCAβ complements a Setaria RCAβ null mutant

To express AtRCA in *Setaria*, we created the transgene shown in Fig. 2A, where the region corresponding to mature AtRCA was preceded by the maize ubiquitin promoter and chloroplast transit peptide. Wild-type *Setaria* Me034 was transformed with this construct, and multiple events were obtained that are referred to as AtRCA in this manuscript, and express both endogenous SvRCA and AtRCA. Two events were advanced to the T2 generation and crossed to SvRCAβ null mutants. Progeny from these crosses yielded AtRCA1[bKO]-1 and AtRCA2[bKO]-2. The experiments in the main text used AtRCA1 and AtRCA1[bKO]-1, simplified to AtRCA and AtRCA[bKO]. Experiments with this line were selectively corroborated using AtRCA2 and AtRCA2[bKO]-2 (see Methods and Results). Transcription of native SvRCA and the AtRCA transgene were assessed for AtRCA and AtRCA[bKO] compared to WT by RT-qPCR (Fig. S2C). This confirmed that native SvRCA was expressed in the WT and AtRCA lines, but not the AtRCA[bKO] line, while the AtRCA transgene was equally expressed in AtRCA and AtRCA[bKO]. WT, AtRCA and AtRCA[bKO] plants were grown at 25°C and ambient CO_2_, showing that addition of the AtRCA transgene to bKO can recover growth and remove the need for 9,000 ppm CO_2_ (Fig. 2B). Overall, these three plant genotypes were similar in growth and appearance, which was verified by statistical analysis of height and tiller count at various ages, as well as dry weight at maturity (Table 1 and Fig. S6). This shows that neither addition of AtRCA to WT *Setaria* nor substitution of AtRCA for SvRCA alters the growth parameters measured when plants were grown at 25°C. Immunoblot analysis revealed similar accumulation of RCA in all 3 lines (Fig. 2C), confirmed by quantitative analysis of multiple lines from both events (Figs. S3C-D and S7), indicating that addition of the AtRCA transgene did not proportionately increase RCA content. This may reflect an endogenous post-transcriptional mechanism that limits RCA accumulation. Because immunoblots cannot distinguish AtRCA from endogenous SvRCA, the proportion of RCA in the AtRCA lines contributed by the AtRCA transgene versus the native SvRCA gene cannot be determined. Given the similarities in transcript levels between AtRCA and AtRCA[bKO] lines, it is possible that AtRCA makes up a substantial proportion of the RCA population in AtRCA lines.

**Fig. 2.**
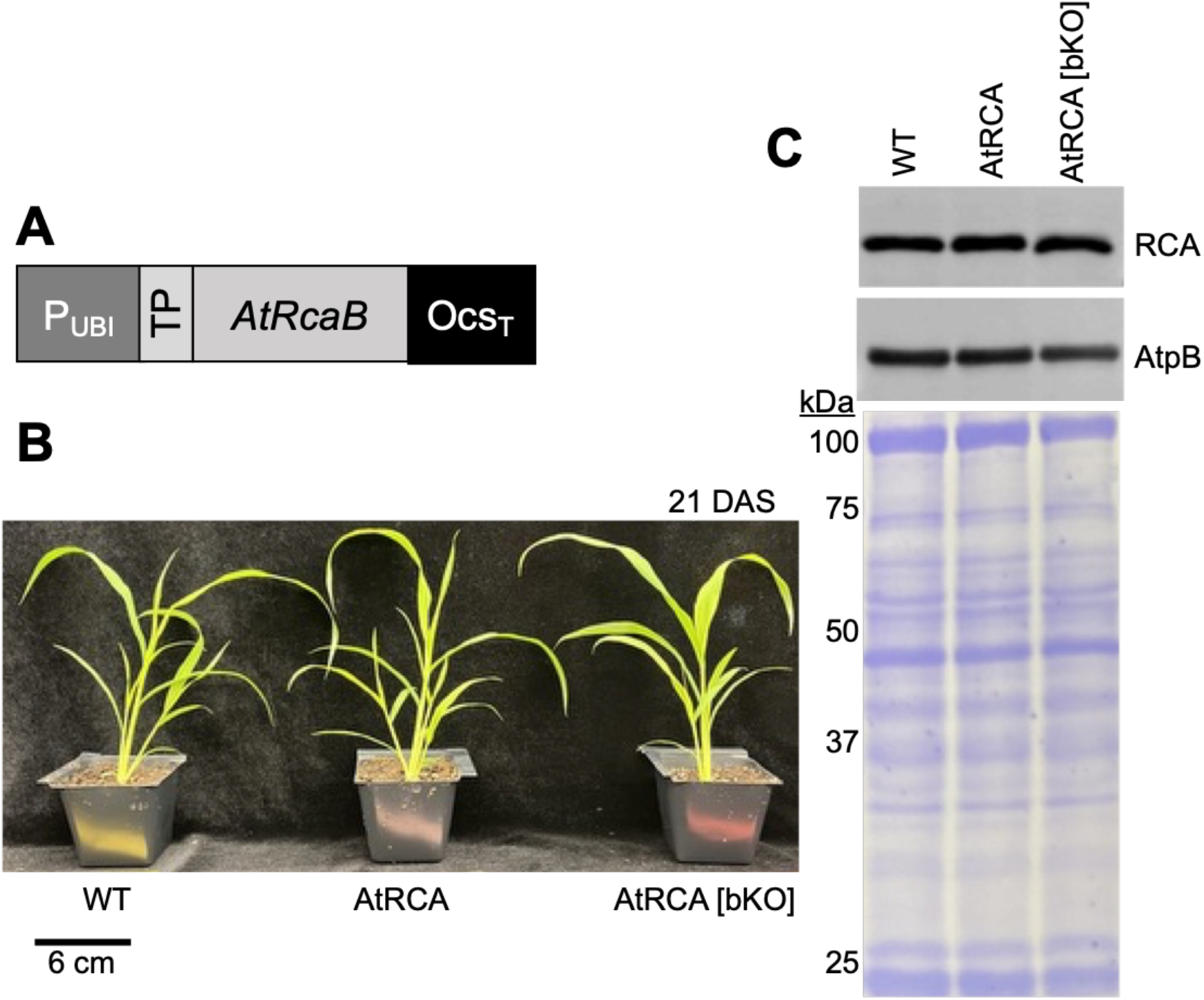
*Agave tequilana* (At) β Rubisco activase (RCA) expression in *Setaria viridis* with (AtRCA) and without (AtRCA [bKO]) *Setaria viridis* wild type (WT) RCAβ. (A) Model of the Agave βRCA (AtRCA) transgene used in this study. P_UBI_, maize ubiquitin promoter; TP, maize chloroplast transit peptide; *AtRcaB*, Agave βRCA coding region; Ocs_T_, octopine synthase terminator. (B) Representative plants at 21 DAS. Plants were germinated on media and transferred to soil after 7 days. (C) Immunoblot analysis of protein isolated from the indicated lines. Antibodies used are shown on the right and Coomassie brilliant blue) stained membrane is below to reflect loading.

**Table 1.**
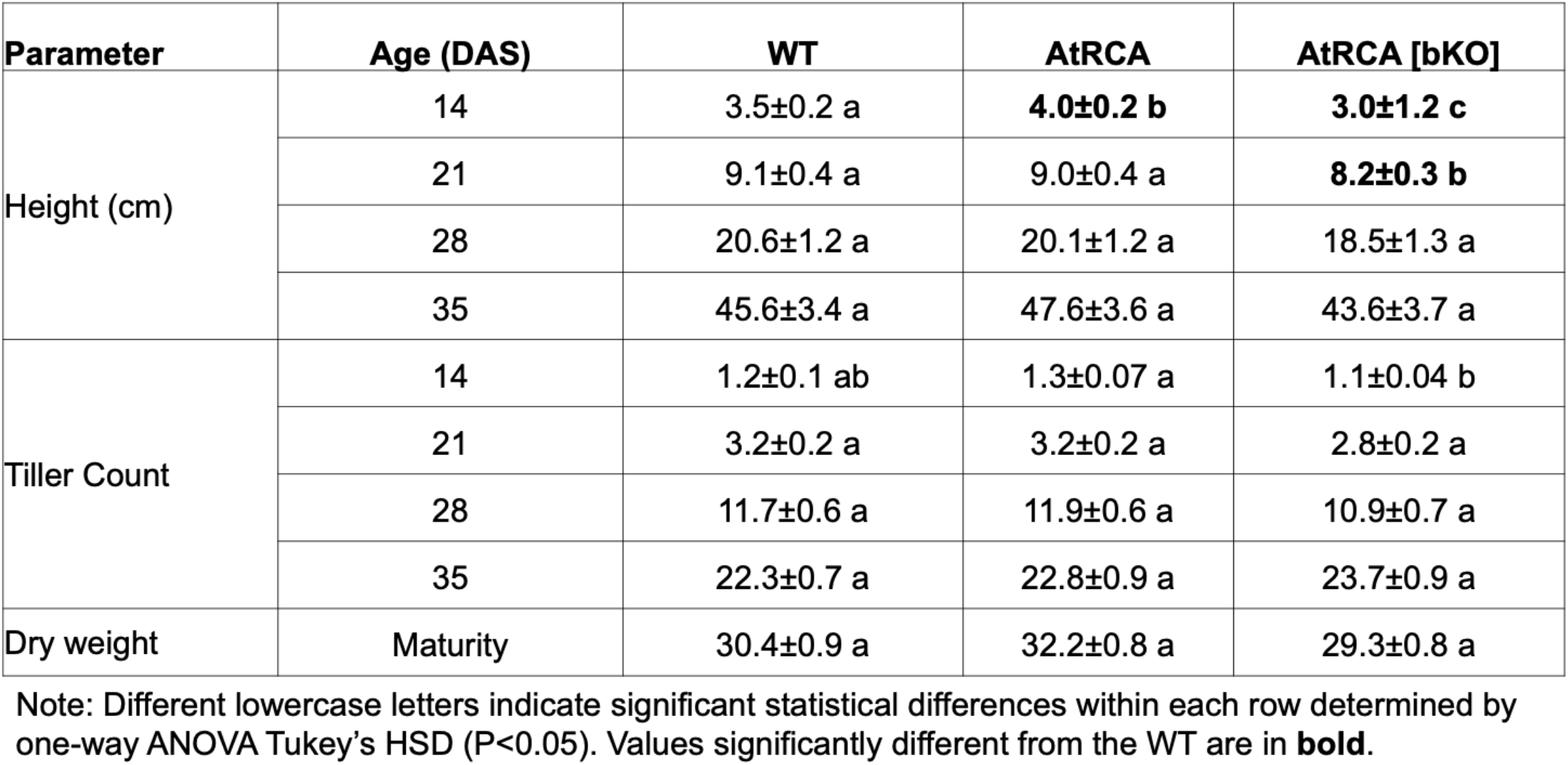
Phenotypic analysis at 35 DAS of WT and transgenic lines grown at 25℃.

### AtRCAβ localizes to both mesophyll and bundle sheath cells

The AtRCA transgene was designed utilizing a ubiquitin promoter to drive expression of AtRCA which is reliable but not cell type specific. *S. viridis* is typical of C_4_ grasses in having Kranz anatomy with mesophyll (M) cells outnumbering bundle sheath (BS) cells, but with BS cells typically having more chloroplasts (Pak *et al*., 1997). While we expected AtRCAβ to accumulate in both M and BS cells, only RCA expressed in BS cells contributes to photosynthesis. To evaluate the localization of AtRCA in the transgenic lines, we separated BS and M cells and assessed protein accumulation in each cell type using immunoblots (Fig. 3). RCA was present in total protein (TP) as well as BS and M preparations for all genotypes (top row). Pyruvate phosphate dikinase (PPDK), used as a marker for M cells, and Rubisco large subunit (LS), used as a marker for BS cells, immunoblots suggested that BS preparations were mostly free of M material, whereas M preparations had some degree of BS contamination.

**Fig. 3.**
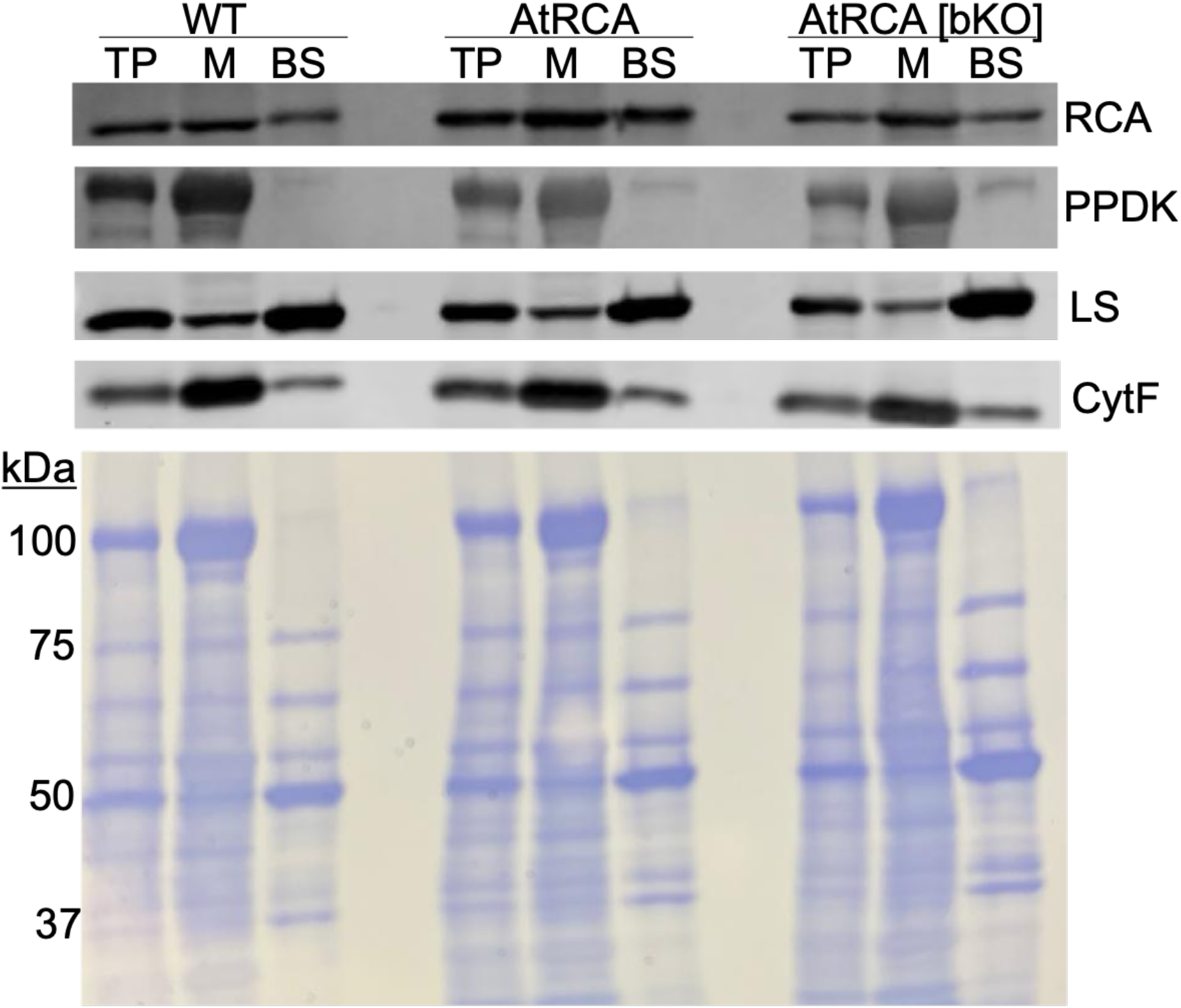
Cell type specificity of proteins in WT and transgenic lines. Immunoblot analysis of proteins isolated from total leaf tissue (TP), mesophyll (M) or bundle sheath (BS) extracts taken from fully expanded leaves of plants 28 DAS. Samples were loaded on an equal protein basis, separated using 8% SDS-PAGE, and membranes were probed with the primary antibodies indicated at right (PPDK, pyruvate, phosphate dikinase; LS, Rubisco large subunit; CytF, cytochrome f). A stained membrane is below to reflect loading.

Quantification of the protein distribution from multiple gels similar to the one shown in Fig. 3 revealed that the M signal for RCA is higher in AtRCA and AtRCA[bKO] than it is in WT. This indicates that some AtRCA is accumulating in M cells, presumably chloroplasts. Nonetheless, the amount of RCA in BS preparations is similar (AtRCA[bKO]) or higher (AtRCA) than WT, leading us to conclude that the overall strength of the ubiquitin promoter is sufficient to restore at least WT RCA protein levels to BS cells. Whether RCA is fully cell-type specific in WT plants is in fact ambiguous: fractionation experiments in a maize proteomic study showed a BS preference but not specificity (Friso *et al*., 2010), a conclusion also supported by our data.

### AtRCA has increased carbon assimilation when grown at 25°C

To test the impact on photosynthesis of adding a foreign RCA to *Setaria*, we measured CO_2_ assimilation in plants grown at 25°C under saturating light (*A*_sat_) and 400 ppm CO_2_, and at saturating CO_2_ and light (*A*_max_) for the WT and two transgenic lines (Fig. 4A). Statistical analysis showed that while WT and AtRCA assimilation were indistinguishable for both parameters, AtRCA[bKO] was approximately 10% higher. While modest, this increase is consistent with the strong ability of AtRCA to activate *Setaria* Rubisco (Fig. 1C). Analysis of CO_2_ assimilation in AtRCA2 and AtRCA2[bKO]2, however, did not show any difference compared to WT (Fig. S8), which could be due to insertional differences between transgenes.

**Fig. 4.**
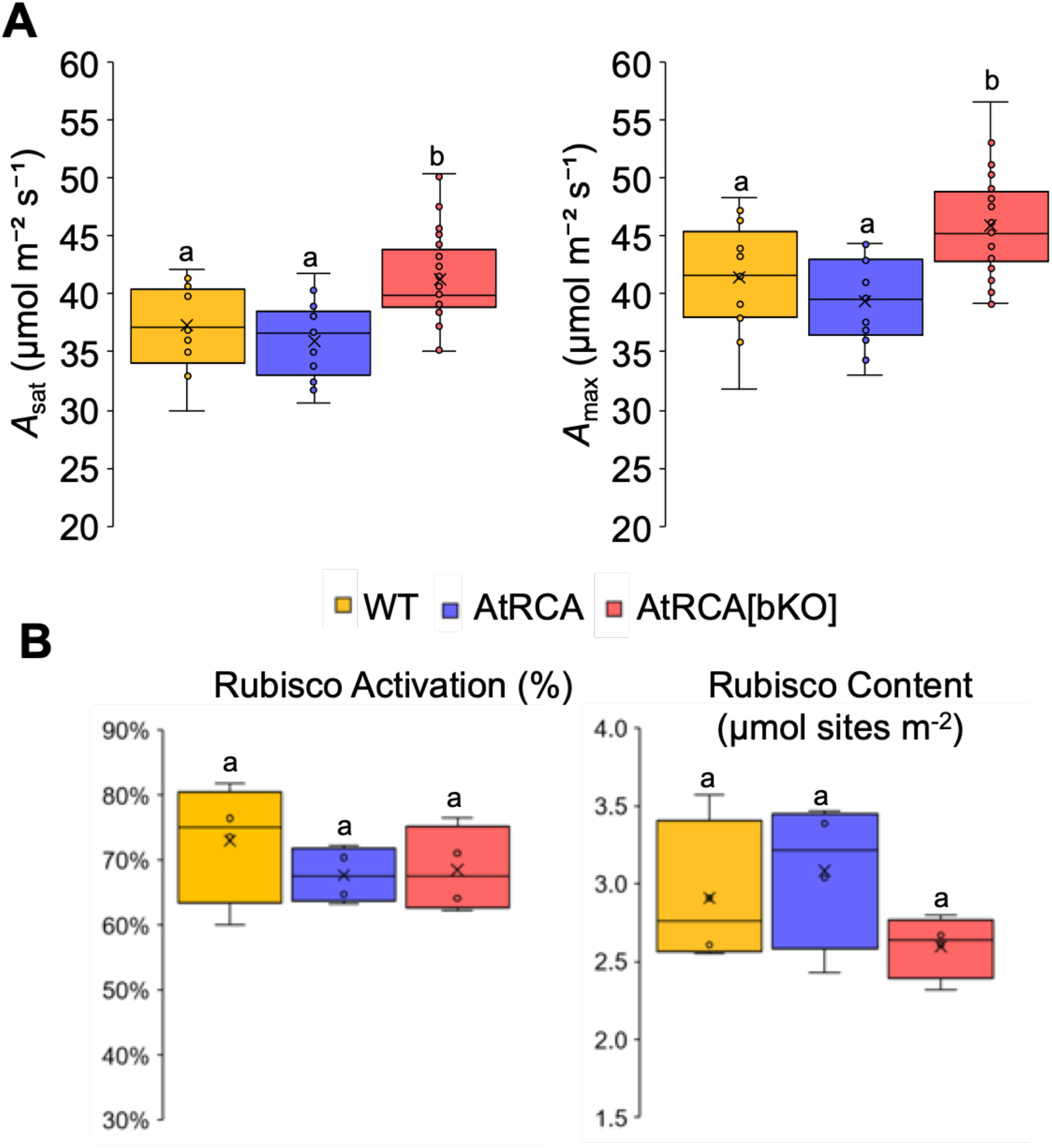
CO_2_ assimilation in transgenic lines compared with the WT. (A) CO_2_ assimilation at saturating light (A_sat_; 1500 µmol m^-2^ s^-1^) and 400 ppm CO_2_, and maximum CO_2_ and saturating light conditions (A_max_). Box plots show the median and first and third quartiles, and the whiskers represent the range. An x represents the mean, and the circles represent individual samples (*A*_sat_ n=16 for WT, 11 for AtRCA, and 27 for AtRCA [bKO]; *A*_max_ n=17 for WT, 11 for AtRCA, and 29 for AtRCA [bKO]). (B) Rubisco activation (left) and content (right) in WT, AtRCA and AtRCA [bKO]. Leaf samples were collected from plants 21 DAS following equilibration at saturating light conditions (1500 PPFD) and 400 ppm CO_2_ to reflect the condition under which *A_sat_* data was collected (n=4). Different lowercase letters show significant differences (P<0.05) determined by one-way ANOVA with Tukey’s HSD test.

To see if the results for *A*_sat_/*A*_max_ could be rationalized by Rubisco parameters, we measured the Rubisco activation state and content from the same plants. Fig. 4B shows that the results for all lines were comparable and thus do not readily explain the higher photosynthetic rate in AtRCA[bKO]. It is possible that Rubisco quantification and activation assays may not easily distinguish a ∼10% difference between samples or that the number of samples analyzed was too small to perceive slight biological differences. Regardless, the increased carbon assimilation in AtRCA[bKO] plants at 25°C did not impact the plant phenotype as measured in this study (Table 1), suggesting that this may not be sustained or significant through the life cycle.

### Carbon assimilation during short-term heat treatment is similar between genotypes

Given the slight increase in photosynthetic rate for AtRCA at control temperatures, and the slightly higher thermostability of AtRCA *in vitro*, we tested the response of photosynthesis to heat treatment in the WT and transgenic lines. We used 44°C as the heat condition, which was higher than the 40°C we previously used as *Setaria* photosynthesis was largely unaffected at that temperature, and *in vitro* data suggests the thermostability of SvRCA and AtRCA ATPase activity may differ at 44°C (Hotto *et al*., 2025). Photosynthesis in *Setaria* was also recently shown to tolerate 42°C for an extended period (Zhang *et al*., 2025).

As shown in Fig. 5, after 6 hr of heat treatment both *A*_sat_ and *A*_max_ showed a similar trend as th e control conditions, with AtRCA[bKO] having significantly higher CO_2_ assimilation than AtRCA and a marginally higher rate than the WT. After 24 hr of heat treatment, assimilation rates for all lines declined by about 50% and were indistinguishable. This suggests that by 24 hr, RCA activity was not limiting photosynthetic carbon assimilation. CO_2_ assimilation was also measured in the second AtRCA event, with and without SvRCA, however, the trends were slightly different. In this case, no differences were observed at the 6 hr time point, but the AtRCA2[bKO]2 line had a significantly higher rate of CO_2_ assimilation after 24 hr compared to AtRCA2 and a minor increase compared to WT (Fig. S8). While results for AtRCA[bKO] and AtRCA2[bKO]2 are slightly divergent, they both suggest that AtRCA may be mildly beneficial to photosynthesis in *Setaria* under heat stress.

**Fig. 5.**
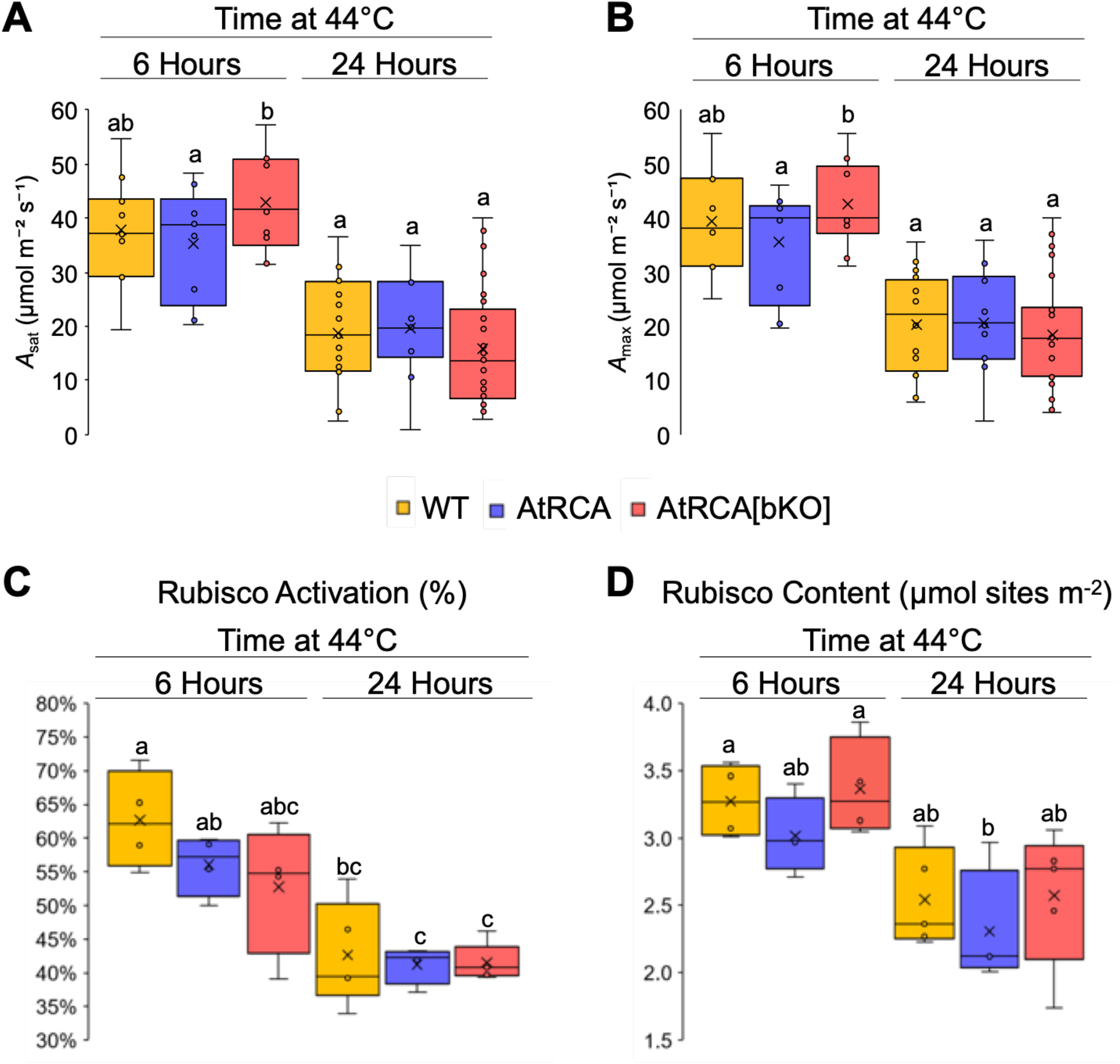
Analysis of photosynthesis parameters in WT and transgenic lines after 6 or 24 hrs at 44°C. (A) CO_2_ assimilation at saturating light (1500 µmol m^-2^ s^-1^) and ambient CO_2_ (*A*_sat_). (B) CO_2_ assimilation at saturating light and maximum CO_2_ (*A*_max_). *A*_sat_ and *A*_max_ n=11 for WT, 9 for AtRCA, and 10 for AtRCA [bKO] at 6 h and 15-16 for WT, 10 for AtRCA and 25-17 for AtRCA [bKO] at 24 h. (C) Rubisco activation and (D) Rubisco content were measured in samples collected following equilibration at saturating light conditions (1500 PPFD) and 400 ppm CO_2_ to reflect the condition under which *A_sat_* data was collected (n=4-5). Plants were grown at 25°C for 25-27 days, after which they were placed at 44°C and CO_2_ assimilation was measured after 6 h and 24 h. Different lowercase letters show significant differences (P<0.05) determined by one-way ANOVA (A and B) or two-way ANOVA (C and D) with Tukey’s HSD test.

To evaluate RCA abundance at 44°C, we analyzed protein extracts by immunoblot (Fig. 6). As expected (Kim *et al*., 2021; Hotto *et al*., 2025), SvRCAα was induced as early as 2 hr, and its accumulation remained stable over the 24 hr time course. RCAβ remained the predominant RCA isoform, either similar in abundance (WT) or increasing modestly in the transgenic lines (Fig. S8) after 24 hr. Since RCAα alone can partially support photosynthesis under heat stress (Hotto *et al*., 2025), the assimilation rates in Fig. 5 reflect activity of SvRCAα combined with either SvRCAβ (WT), AtRCAβ (AtRCA[bKO]) or both (AtRCA).

**Fig. 6.**
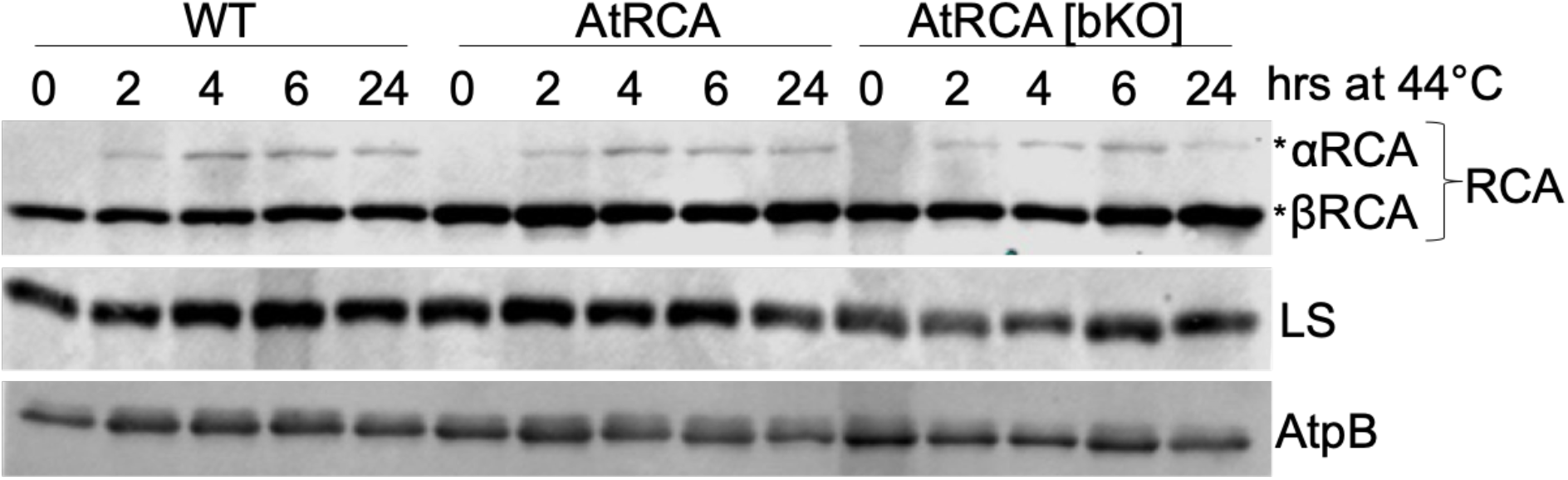
Induction of αRCA and stability of βRCA at 44°C in transgenic lines and the WT. Immunoblot analysis of total protein isolated from the youngest fully expanded leaf from plants grown at 25°C for 25-27 days, then transferred to 44°C for the time indicated. Samples were separated by 8% SDS-PAGE, loaded on an equal leaf area basis, and blots were probed with the antibodies indicated at right.

To determine whether Rubisco activation state responded to heat treatment, we analyzed leaf tissue after 6 and 24 hr at 44°C (Fig. 5C). Whereas the activation state had been 70-75% at 25°C (Fig. 4B), after 6 hr at 44°C the activation state had declined to 55%-62%, with a further, significant, decline to 40-45% after 24 hr. Thus, Rubisco activation was more sensitive to heat *in vivo* than the *in vitro* Rubisco activation potential or ATPase activity of RCA (Figs. 1B and 1D). This suggests that RCA stability and activity may not be the limiting factor for Rubisco activation at higher temperatures, or the thermostability of RCA *in vivo* is distinct from its *in vitro* thermostability.

Changes in Rubisco content could also impact CO_2_ assimilation in response to heat. Measurements showed that after 6 hr at 44°C Rubisco content was similar or slightly higher (Fig. 5D) than what was measured from plants at 25°C (Fig. 4B). After 24 hr, Rubisco content returned to pre-stress amounts in all lines. A heat-induced decline in Rubisco content has been previously reported in other C_4_ species such as maize (Perdomo *et al*., 2017), which is exacerbated by other cellular factors that diminish Rubisco catalysis (Crafts-Brandner and Salvucci, 2002). The response of Rubisco content to growth at 44°C in *Setaria* has not been previously reported. Overall, we conclude that the Rubisco activation state limits photosynthetic performance in all lines after 24 hr at 44°C.

### Dynamic responses of photosynthesis are similar in all lines

We used two protocols to test whether the photosynthetic rates differ between lines under dynamic conditions, thereby querying short-term responses rather than longer term metabolic adjustments. We first used a heat curve, where the temperature of a clamped leaf was increased by 2°C every 10 min, with light set to a medium intensity of 400 µE to avoid photoinhibition. Fig. 7 shows that while the values for individual plants varied, for all lines the highest photosynthetic rate was attained at 38-40°C, followed by a decline to the highest tested temperature of 54°C. Surprisingly, AtRCA[bKO] had the highest photosynthetic rate at 38°C compared to 40°C for both WT and AtRCA, although these differences were not statistically significant. This contrasts with the *in vitro* data that showed AtRCA having a much higher thermostability, with ATPase and Rubisco activation activity declining at temperatures greater than 45°C. An important caveat is that *in vitro*, only RCA is heated whereas whole-plant assays heat the entire system. It should also be noted that in this experiment, the time duration of the heat treatment is likely too short to reflect induction of SvRCAα, although this was not measured. The shapes of the heat curves were also similar between genotypes, suggesting a similar dynamic response to changing temperature. Overall, the near overlap of the WT and AtRCA[bKO] lines suggests that there is no effect of the RCA type on the heat response measured here.

**Fig. 7.**
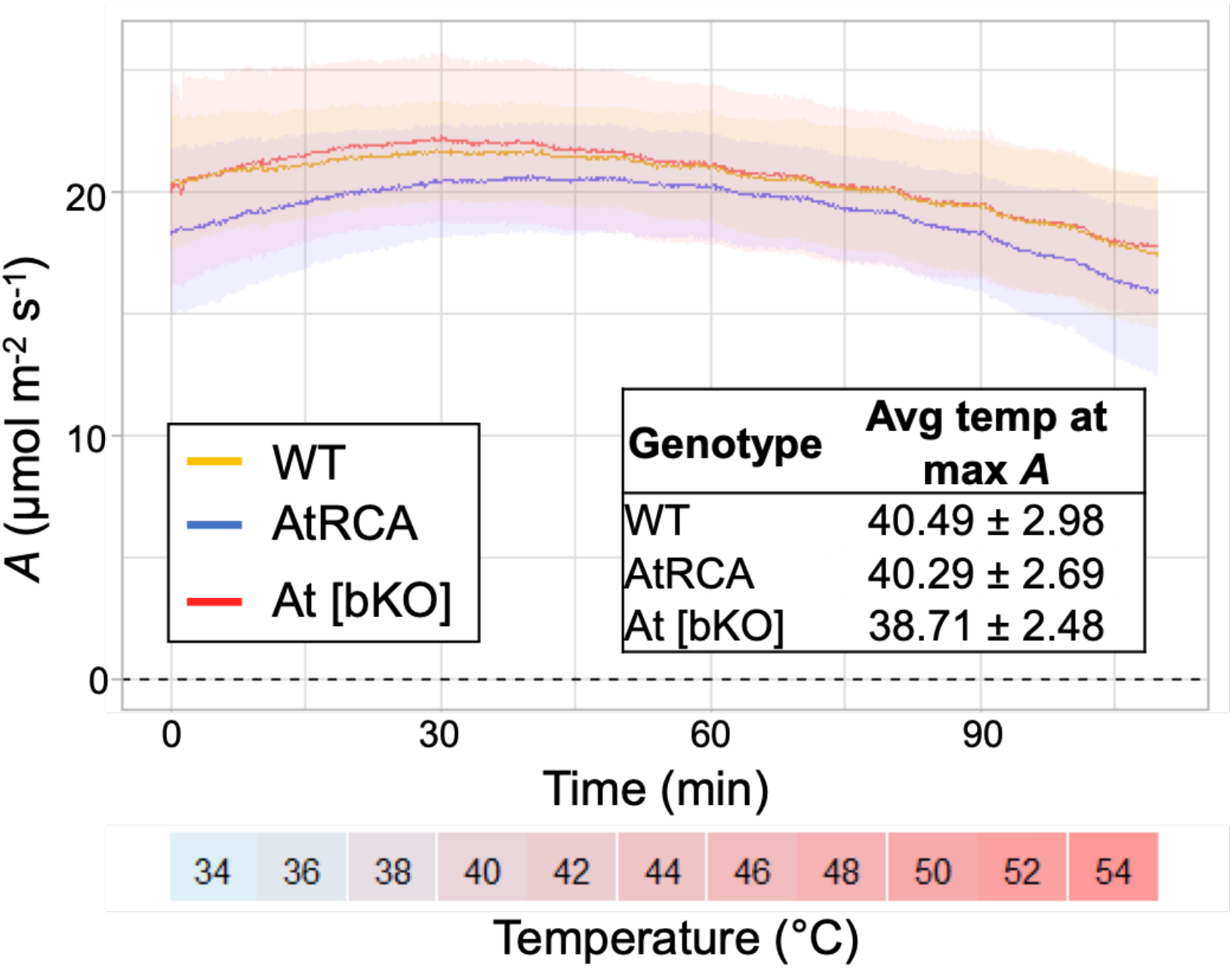
CO_2_ assimilation between 34°C and 54°C. An LI-6800 was used to measure the response of CO_2_ assimilation rates to increasing temperatures from the youngest fully expanded leaf on the primary tiller at 25-29 DAS. The temperature of the LI-6800 head increased by 2°C every 10 minutes and the light conditions were set to 400 PPFD. The shading around each line represents the standard deviation for each genotype (n=9 for WT and AtRCA, and n=8 for AtRCA [bKO]).

A second protocol measured the recovery of photosynthesis after dark adaptation, a process that is limited in part by the light-stimulated reactivation of Rubisco in some species (Wang *et al*., 2021). For these experiments, a leaf was dark adapted for 30 min, then photosynthesis was measured following the initiation of 1500 µE light using an LI-6800. Fig. 8 shows that when the responses were normalized to the peak photosynthetic rate achieved after 30 min of light, kinetics varied little between the WT and transgenic lines. All lines had similar photosynthesis induction rates within the first few minutes, and AtRCA[bKO] containing only AtRCA lagged somewhat in the second phase to reach maximum photosynthetic capacity. These results suggest that SvRCA and AtRCA are equivalently effective in helping to support light induction of photosynthesis under the tested conditions.

**Fig. 8.**
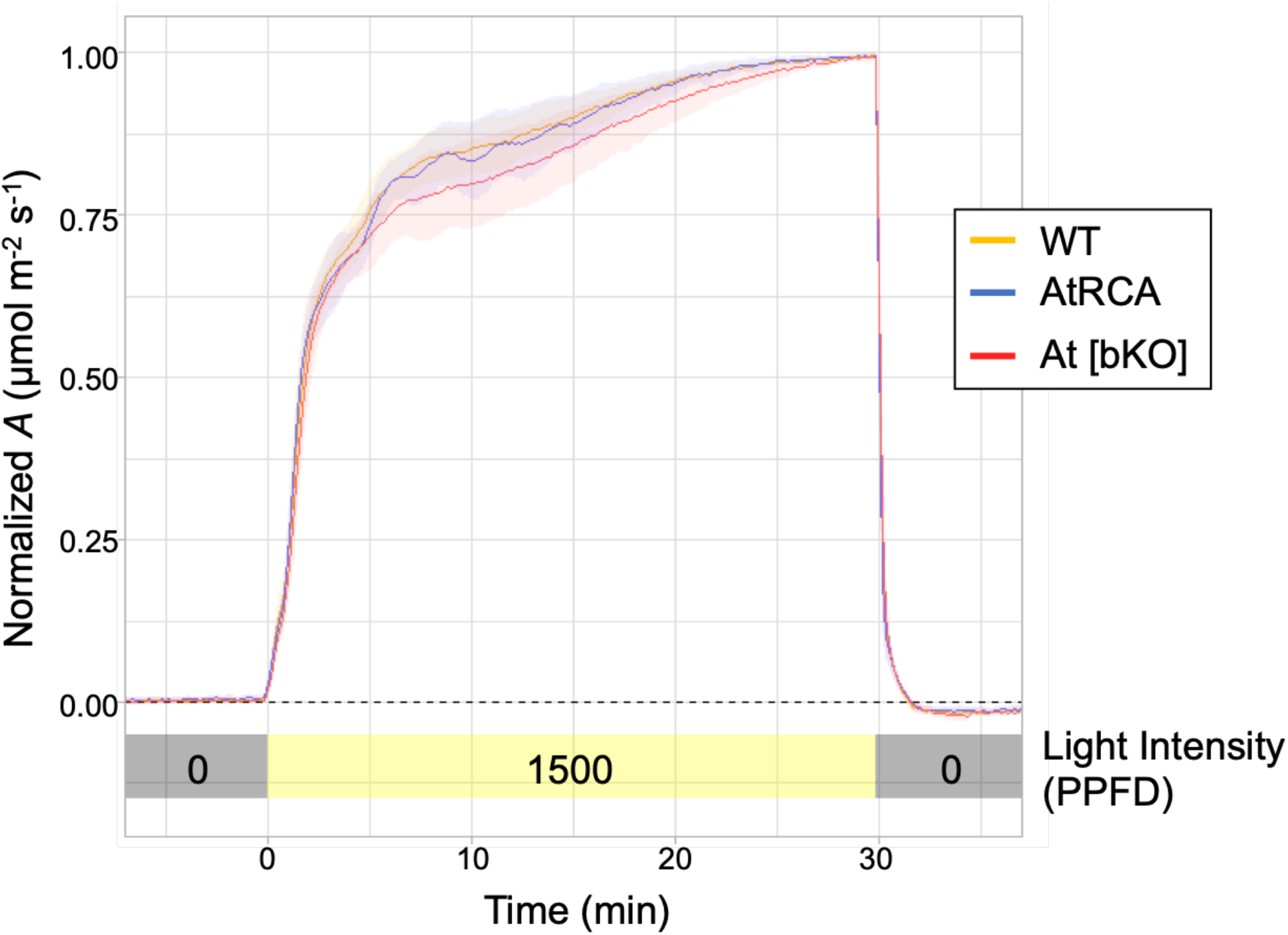
CO_2_ assimilation in response to light fluctuation. Response of CO_2_ assimilation rates to saturating light (1500 PPFD) at 25°C after 30 minutes of dark acclimation (0 PPFD) measured using an LI-6800 from the youngest fully expanded leaf on the primary tiller at 25-29 DAS. The shading around each line represents the standard deviation for each genotype (n=14 for WT, 15 for AtRCA and 13 for AtRCA [bKO]).

## Discussion

Stabilizing photosynthesis under increased temperatures has long been a target for crop improvement, especially in an era where the mounting frequency and duration of heat stress are increasing pressure on agriculture (Mahajan *et al*., 2025). Given the propensity for Rubisco deactivation at increased temperatures and the variation of RCA thermotolerance, improving Rubisco activation under heat through RCA engineering has significant potential (Qu *et al*., 2023), although the possibilities have been little explored in C_4_ plants. Here we have evaluated the properties of *Setaria viridis* plants to which we have added expression of RCAβ from *Agave tequiliana* (line AtRCA) or substituted *Agave* RCA for endogenous RCAβ (line AtRCA[bKO]). We chose *Agave* RCAβ because of its extraordinary thermostability reported *in vitro* (Shivhare and Mueller-Cajar, 2017), reasoning that it might further enhance the photosynthetic heat tolerance of *S. viridis*. *Agave* uses CAM photosynthesis where initially fixed CO_2_ is released into the Calvin cycle in the daytime, and thus Rubisco activity is adapted to high temperatures. Our results demonstrate that a C_4_ RCA can be substituted with a CAM RCA *in planta* despite their evolutionary and biochemical divergence and also points to potential limitations of this approach.

### *In vitro* assays are valuable in the preliminary characterization of diverse Rubisco activases

There have been limited *in vitro* studies of recombinant RCAs from C_4_ plants and the single CAM RCA from *Agave*, in contrast to extensive biochemical investigations of C_3_ activases that have established the majority of RCA core properties (Bhat *et al*., 2017; Waheeda *et al*., 2023). Detailed analysis of *Agave* RCA demonstrated that its thermotolerance maps to the Rubisco activation domain, and that the protein remains soluble under conditions where rice RCA denatures (Shivhare and Mueller-Cajar, 2017). These characteristics, along with a demonstration of high activity on distantly related rice Rubisco, made AtRCA an attractive candidate for engineering into *Setaria*. Other potential but as yet unexplored sources of exceptionally thermotolerant RCAs include the C_4_ desert plant *Tidestromia oblongifolia* and even thermophilic cyanobacteria (Ogbaga *et al*., 2018; Prado *et al*., 2025).

Prior to evaluation of transgenic plants, we verified that AtRCAβ was able to activate *Setaria* Rubisco, since neither *in vivo* substitution of C_4_ RCAs nor transgenic introduction of a CAM RCA into a C_4_ plant had been reported. A side-by-side comparison showed a gradient of thermotolerance of ATPase activity with AtRCAβ the most thermotolerant, followed by SvRCAβ and ZmRCAβ, the most thermosensitive of the RCAs assayed in this study (Fig. 1). The properties of SvRCAβ were superior to those of the C_3_ desert creosote *Larrea* RCA that lost ATPase activity and Rubisco activation activity at temperatures exceeding 35°C and 40°C, respectively (Salvucci and Crafts-Brandner, 2004). This suggests that select C_4_ RCAs may harbor traits lending thermostability that have yet to be explored in detail.

ATPase activity of the recombinant RCAs did not necessarily correlate with their Rubisco activation efficiency at the same temperatures. Previous *in vitro* analysis of AtRCA in activating rice Rubisco showed a similar trend – a lower ATPase activity but a higher Rubisco activation ability (Shivhare and Mueller-Cajar, 2017). In the same study, rice RCAβ had a higher ATPase activity and lower Rubisco activation ability, which was the trend we observed with SvRCA in activating the cognate Rubisco. Additionally, all three RCAs assayed here had higher thermotolerance for Rubisco activation compared to ATPase activity, suggesting that Rubisco activation is not solely determined by ATPase activity. Overall, these results confirmed that AtRCA was effective at activating *Setaria* Rubisco and had a higher thermotolerance than SvRCAβ.

As AtRCA plants are presumed to express both AtRCA and SvRCA, we examined the *in vitro* properties of 1:1 mixtures of the two isomers (Figs. 1 and S4). While the properties of mixtures were intermediate and thus consistent with heterooligomerization, we cannot formally differentiate between the presence of heterooligomers and a mixture of homooligomers. Because of the cooperativity of RCA subunits, previous studies have used an RCA inactivated through an ATPase active site mutation to show its disproportionate effect on the activity of hexamers (Van De Loo and Salvucci, 1998). Other studies showed that heterooligomerization can be temperature-dependent, with a heat-sensitive isoform becoming insoluble and thus excluded from hexamers at elevated temperature (Scafaro *et al*., 2019). A similar result was obtained with mixtures of *Agave* and rice RCA, with only *Agave* remaining soluble at higher temperatures (Shivhare and Mueller-Cajar, 2017). Thus, it is possible that heat-treated AtRCA plants experience a change in RCA hexamer composition as compared to plants grown at 25°C.

While we did not carry out a detailed study of interactions between AtRCA and SvRCA, we performed a preliminary experiment using a D173A mutation in SvRCA that inactivates ATPase activity as first shown for the tobacco enzyme by mutations at D174 (Van De Loo and Salvucci, 1998). Recombinant D173A was indeed inactive as an ATPase and thus had no activity on purified Rubisco (Fig. S9A). 1:1 mixtures of SvRCA and D173A resulted in a decrease in Rubisco activation by about 70%, consistent with cooperativity (Fig. S9B). A similar decrease was observed with mixtures of AtRCA and D173A, which suggests that AtRCA and SvRCA also freely heterooligomerize. This test is relevant although sometimes lacking when interpreting the phenotypes of transgenic lines expressing RCA from multiple species which are inevitably exposed to heat stress, as has been done extensively in rice through addition of maize RCA (Fukayama *et al*., 2012; Qu *et al*., 2021; Yamori *et al*., 2026), as well as reciprocally (Feng *et al*., 2023). Arabidopsis has also been used as a foreign RCA recipient, but in contrast the variants are typically introduced into the *rca* null mutant background, akin to what we have shown here (Kurek *et al*., 2007; Kumar *et al*., 2009; Carmo-Silva and Salvucci, 2013).

### A CAM RCA can replace a C_4_ RCA in Setaria

Using the maize ubiquitin promoter, we were able to achieve WT-like levels of AtRCA expression as most clearly evidenced by RCA accumulation in AtRCA[bKO] lacking endogenous RCAβ. In the primary transformants expressing both SvRCA and AtRCA, overall RCA accumulation was similar to WT, although in both cases a portion of that RCA was localized to mesophyll cells where it is presumably functionally neutral. Immunoblots and photosynthetic data from the AtRCA[bKO] lines most clearly show the functionality of the AtRCA transgen<u>e</u>. In the AtRCA lines, there appears to be an inherent limitation to RCA accumulation since the amount of RCA is not the sum of WT and AtRCA[bKO]. We can therefore only infer that both SvRCA and AtRCA accumulate in the AtRCA lines, and that the ratios of the two proteins could change dynamically with plant growth conditions.

The ability of RCA to function across species is well documented, with the unique exception of the *Solanaceae* (Wang *et al*., 1992). The most recent common ancestor for *Setaria* and *Agave* being some 125 Mya, however (Wang *et al*., 2024) did raise the possibility of a cross-species incompatibility. Mutational studies of AtRCA led to the suggestion that the low concentration of Rubisco in CAM plants may have led to the evolution of stronger Rubisco binding (Shivhare and Mueller-Cajar, 2017), which would be consistent with our observation of strong activation of *Setaria* Rubisco by AtRCA (Fig. 1) as well as the robust performance of transgenic plants. Even if AtRCA were slightly less active across species *in vivo*, earlier studies showed that downregulation of RCA did not affect steady-state photosynthetic rates until the decrease became rather drastic. This has led to the conclusion that RCA is generally present in excess both in C_4_ (von Caemmerer *et al*., 2005) and C_3_ species (Mate *et al*., 1996; Yamori and von Caemmerer, 2009).

Other studies, mainly in rice, suggest conversely that increasing activase content, at least in C_3_ plants, can augment photosynthesis under certain conditions. As mentioned above, Yamori *et al*. (2026) found that addition of maize RCAβ to rice improved responses to fluctuating light, in contrast to our finding that in light to dark transitions, no difference between genotypes could be observed (Fig. 8). Also, in rice, co-overexpression of RCA and RBCS improved plant performance under heat stress (Qu *et al*., 2021) but not at normal temperatures (Suganami *et al*., 2021). In Arabidopsis, addition of an RCA gene from the halophyte *Suaeda salsa* benefitted photosynthesis under salt stress (Yang *et al*., 2026), while homologous activase overexpression in cucumber was favorable to growth under low temperature and light conditions (Bi *et al*., 2017). Taken together, these and other studies suggest that effects of activase depletion and overexpression are highly contextual, and the transgenic lines we have created could provide additional insights under experimental conditions we have not yet tested.

### At 40°C, addition of AtRCA may confer a slight photosynthetic advantage

One of the main hypotheses of our approach was that the temperature sensitivity of photosynthesis could be altered by adding and/or substituting *Agave* RCAβ for SvRCAβ. *In vitro* assays showed that AtRCA is slightly more thermostable than SvRCA (Fig. 1), and as the plant growth temperature increased, we posited that Rubisco deactivation might be mitigated. Indeed, after 6 hr at 44°C line AtRCA[bKO] had marginally higher *A*_sat_ and *A*_max_ than the other lines (Fig. 5), however this was also true at normal growth temperature (Fig. 4). In addition, endogenous RCAα is induced when the temperature is raised (Fig. 6), confounding data interpretation. For example, one could speculate that if our plants were unable to induce RCAα, a greater differentiation might have been observed between the WT and AtRCA-expressing plants. The close linkage of the *RCAA* and *RCAB* genes, however, precluded crossing the ΔrcaA CRISPR-induced mutation into AtRCA[bKO] lines. A future approach might be to use gene editing to inactivate *RCAA* in the bKO line, creating the potential for a single platform to test exogenous RCA through transgenesis without endogenous RCAα or RCAβ.

After 24 hr at 44°C, assimilation was strongly reduced in all lines as was Rubisco content and activation state (Fig. 5). Considering other C_4_ species, the content decrease was in the range of 20-25%, similar to what was reported for maize grown at 38°C (Perdomo *et al*., 2017), with a greater effect on its activation state. Another study in maize found heat did not affect total Rubisco activity but strongly affected the activation state (Crafts-Brandner and Salvucci, 2002), even well below the *in vitro* Tm of 41.7° that we measured (Fig. 1C). Our prior work where *Setaria* was placed at 40°C and assayed after two or seven days, showed that WT plants did not exhibit a change in assimilation, but may have slightly decreased Rubisco content (Hotto *et al*., 2025). Two other studies found that photosynthetic rates did not decline in *Setaria* grown at 42°C (Stainbrook *et al*., 2024; Zhang *et al*., 2025), albeit in accession A10, which is nonetheless highly similar to accession ME034 used here (Thielen *et al*., 2020). Thus, there may be a breakpoint in temperature sensitivity of photosynthesis in *S. viridis* just above 42°C.

## Conclusions

The results reported here represent a path towards modifying carbon assimilation through engineering of Rubisco activase in a C_4_ platform. Considering that overexpression of maize activase in rice can generate favorable photosynthetic phenotypes (Yamori *et al*., 2026), interchangeability of C_3_, C_4_ and CAM activases may be broadly applicable to plant modification strategies (apart from the Solanaceae; Wang *et al*., 1992). In terms of engineering carbon assimilation, cross-species compatibility of RCA contrasts with the challenges of engineering its client protein Rubisco. The primary reasons are the dual genomic localization of its component subunit genes, and that Rubisco assembly factors tend to be species specific (Qin *et al*., 2025; da Silva *et al*., 2026). Additionally, unlike organisms such as Arabidopsis and *C. reinhardtii* where there is a single RCA gene that can be mutated and replaced by other RCA genes (Kumar *et al*., 2009; Esquivel *et al*., 2013), C_4_ grasses have closely linked duplicated RCA genes, one of which is inducible by heat stress (Nagarajan *et al*., 2025). Given that heat tolerance of photosynthesis is often cited as a rationale for RCA engineering (Ogbaga *et al*., 2018; Sparrow-Muñoz *et al*., 2023), genetic ablation of both genes is a worthy goal for future engineering approaches in species that harbor such duplications.

## Supplementary data

The following supplementary data are available at JXB online:

## Acknowledgements

We thank Oliver Mueller-Cajar (Nanyang Technological Univ.) for the pHUE vector and *Agave tequilana* DNA sequence, Rob Sharwood (Univ. Western Sydney) for the pHUsp2-cc vector, Zhen Guo Oh for RuBP synthesis and clarifying discussions, and Kevin Baudry for advice on statistical analysis and helpful discussions.

## Author contributions

AMH, DBS: Conceptualization and funding acquisition; AMH, DBS: writing; AMH, SG, KE: investigation and formal analysis.

## Data availability

Any data not available in the manuscript and supplementary materials can be requested directly from the authors.

## Abbreviations

RCA: Rubisco activase
WT: wild-type
DAS: days after sowing
PFFD: photosynthetic photon flux density

---

**Fig. S1.**
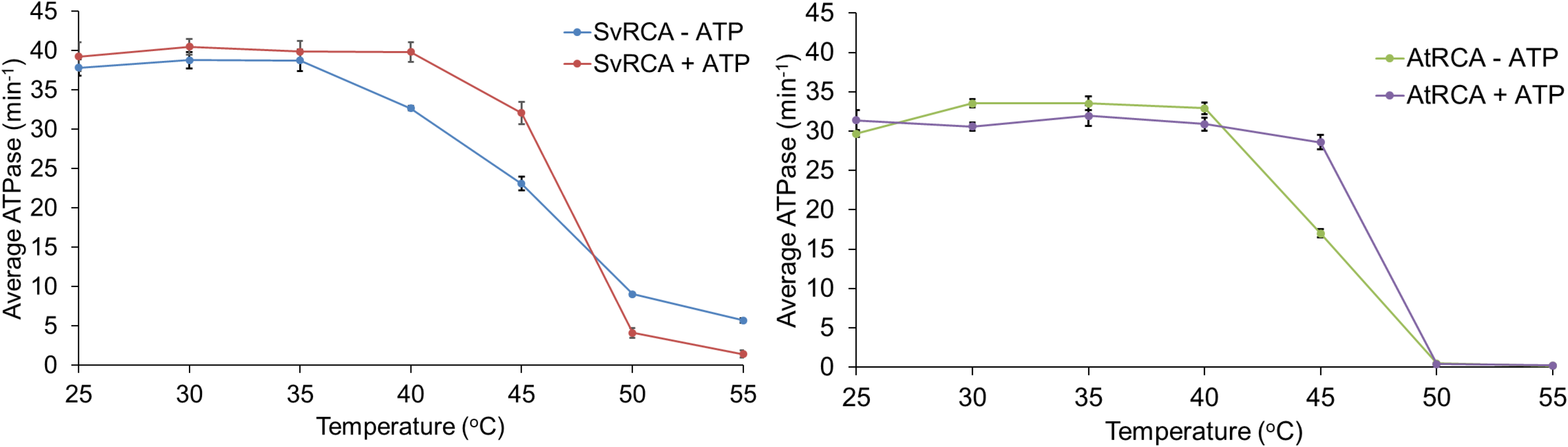
***In vitro* ATPase activity** of Setaria (Sv, left) and Agave (At, right) RCAβ isoforms with and without preincubation for 10 min with 0.2 mM ATP at the indicated temperatures. The ATPase assays were for 10 min at 25°C. Data represent an average ± SE of 2-3 biological replicates with at least 3 technical replicates each.

**Fig. S2.**
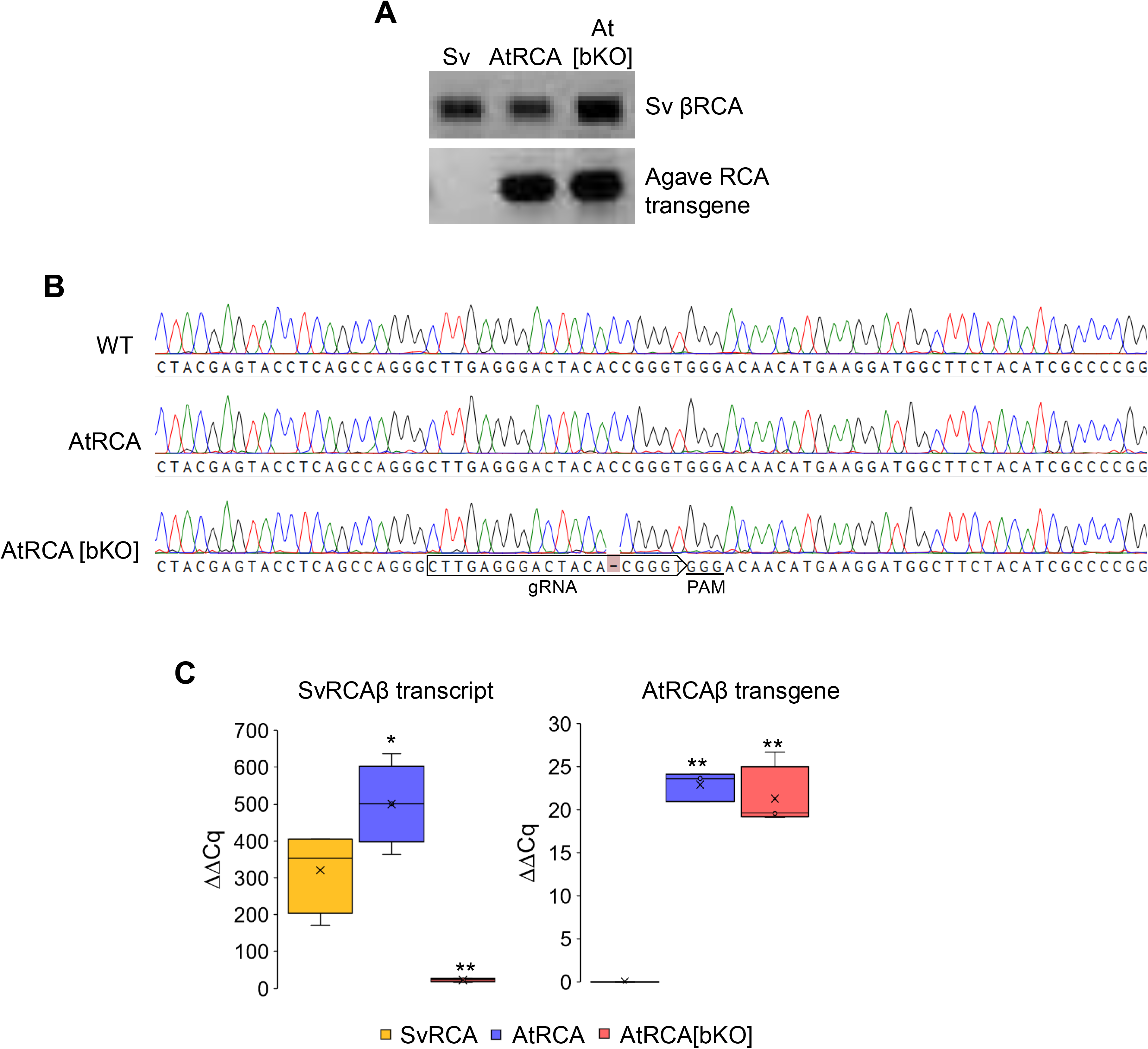
Verification of transgenic lines from AtRCA event 1 used in this study expressing Agave βRCA with (AtRCA) and without (At[bKO]) Setaria βRCA. (A) PCR analysis indicating the presence of the *Setaria* WT and/or AtRCAβ gene. (B) Sequencing results from the *Setaria* WT βRCA gene shows the WT gene present in the WT and AtRCA lines, while the single base deletion creating a premature stop codon in line βKO2 (Hotto *et al*., 2025) is present in AtRCA[bKO]. (C) qRT-PCR analysis of the Setaria βRCA (left) and Agave βRCA (right) transcripts in the three lines used in this study. Significant differences compared to WT calculated from the Student’s *t*-test are indicated (* *p*<0.05; ** *p*<0.001).

**Fig. S3.**
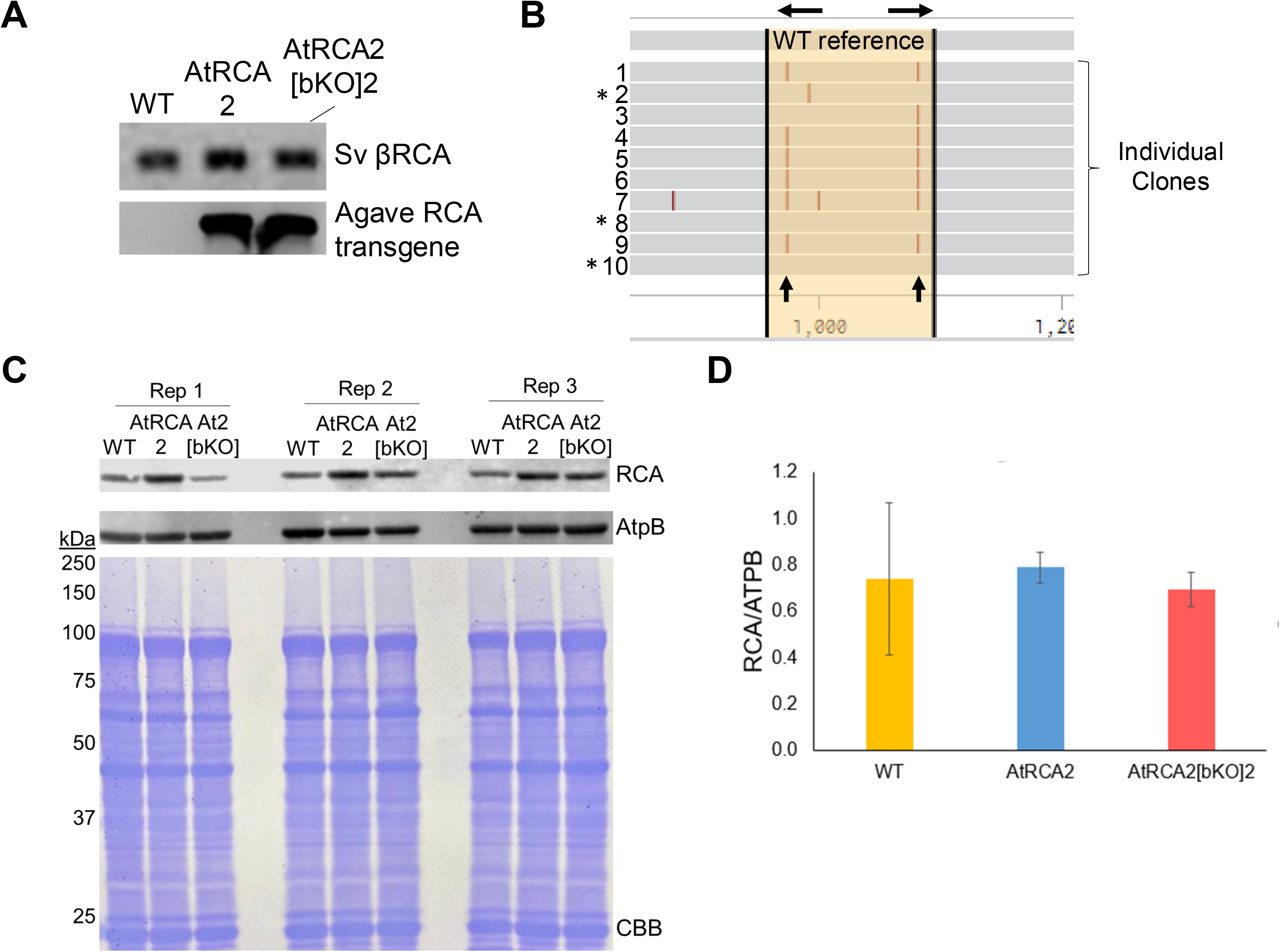
Analysis of a second AtRCA transgenic event expressing Agave βRCA with (AtRCA2) and without (AtRCA2[bKO]2) Setaria βRCA. This line was created from a cross between AtRCA2 and line βKO1 described in Hotto et al. (2025), which contains two single base deletions in contrast to the single deletion in βKO2 (Fig. S2B). (A) PCR analysis of the Setaria WT βRCA gene and the Agave βRCA transgene. (B) Illustration of 10 individual clones sequenced from the Setaria WT βRCA gene amplicon in (A). Black arrows at the top indicate the relative position of gRNAs, while the vertical black arrows at the bottom indicate the positions of anticipated single nucleotide deletions due to gene editing. Three clones with a * represent a WT βRCA amplicon or an amplicon with a silent mutation. (C) Immunoblot analysis of protein isolated from three independent sets of wild type (WT) and Agave complemented lines. Antibodies used are shown on the right and Coomassie brilliant blue (CBB)-stained membrane is below to reflect loading, with molecular mass indicated on the left (kDa; AtpB, ATP synthase β subunit). (D) Quantification of total βRCA relative to AtpB (n=3). Error bars are ± SD.

**Fig. S4.**
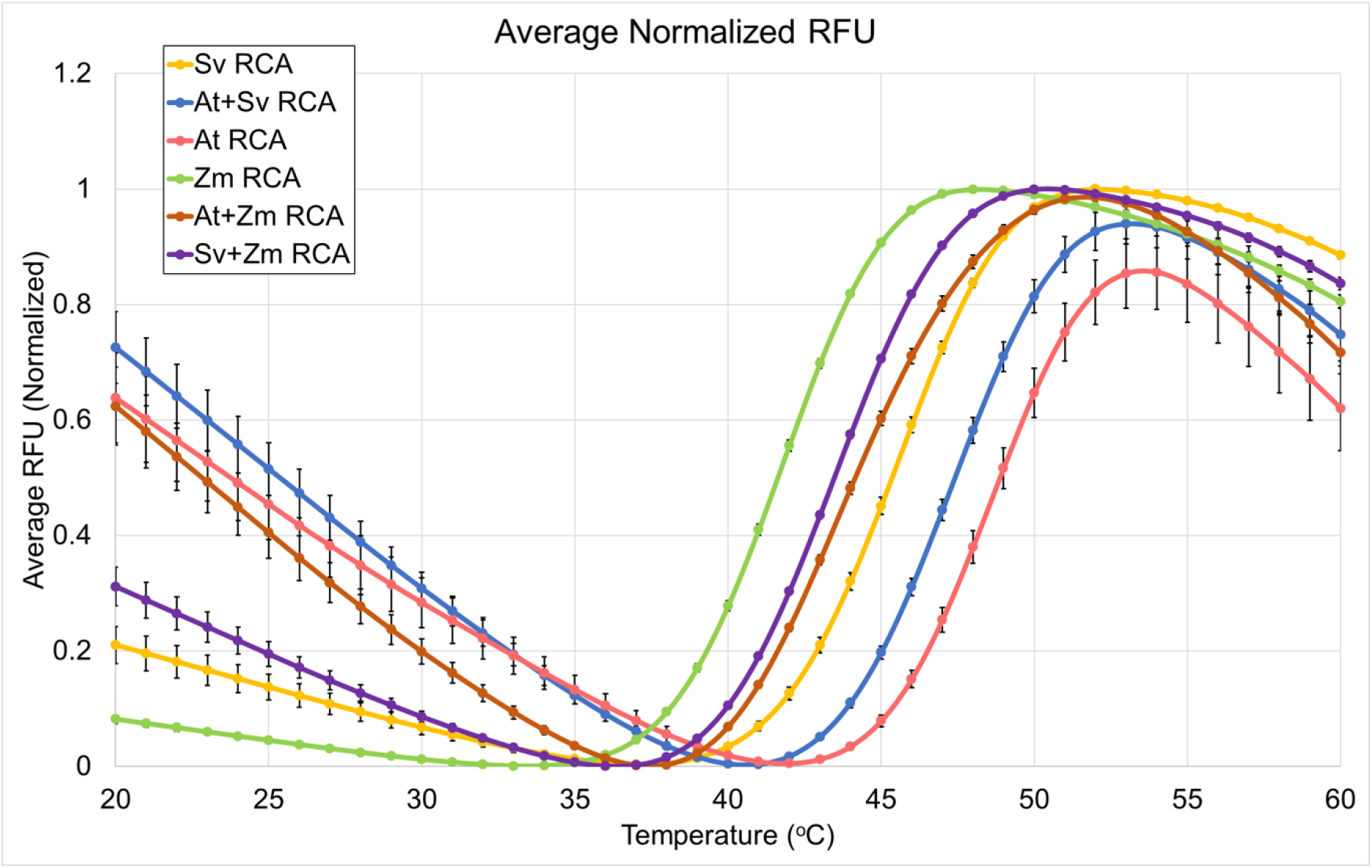
Thermal shift assay of different RCA isoforms singly or in a 1:1 molar ratio. Samples were heated from 20°C to 60°C at 1°C min^-1^. Relative fluorescence units (RFU) were normalized to the maximum fluorescence per sample and sigmoidal curves were fitted to the data. Data represent an average ± SE of 2-3 biological replicates with at least 3 technical replicates each. Sv, *Setaria viridis*; At, *Agave tequilana*; Zm, *Zea mays*.

**Fig. S5.**
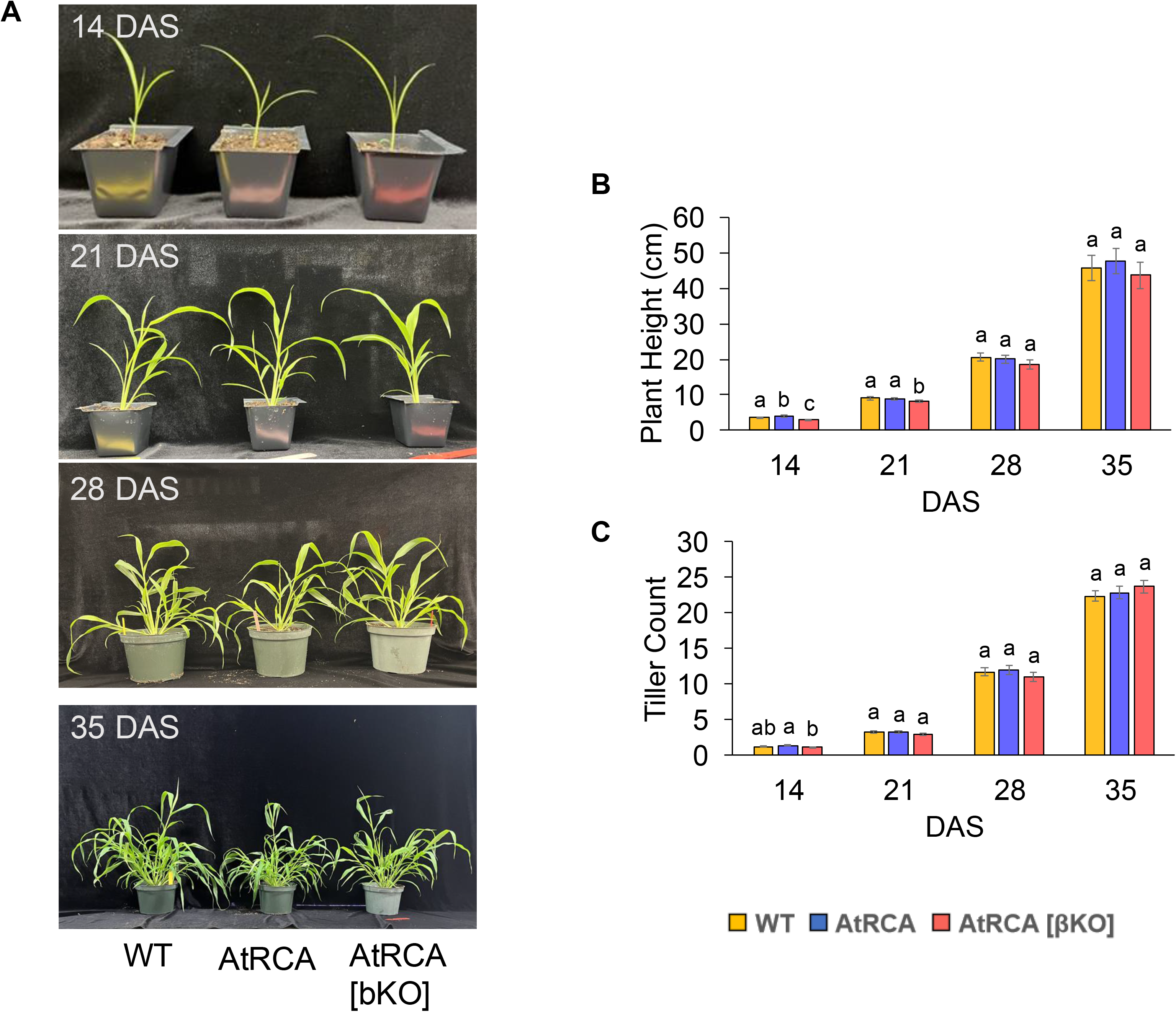
Plant growth at different maturity stages. The WT was compared to the transgenic lines shown (event 1) at 14, 21, 28 and 35 days after sowing (DAS). (A) Representative plant photos taken every 7 days from 14-35 DAS. (B) Plant height was measured from the top of the soil to the base of the youngest fully expanded leaf on the primary tiller (n=18). (C) Tiller count includes all tillers originating from the base of the plant or other tillers (n=18). Different lowercase letters show significant differences (P<0.05) determined by one-way ANOVA with Tukey’s HSD test at each time point.

**Fig. S6.**
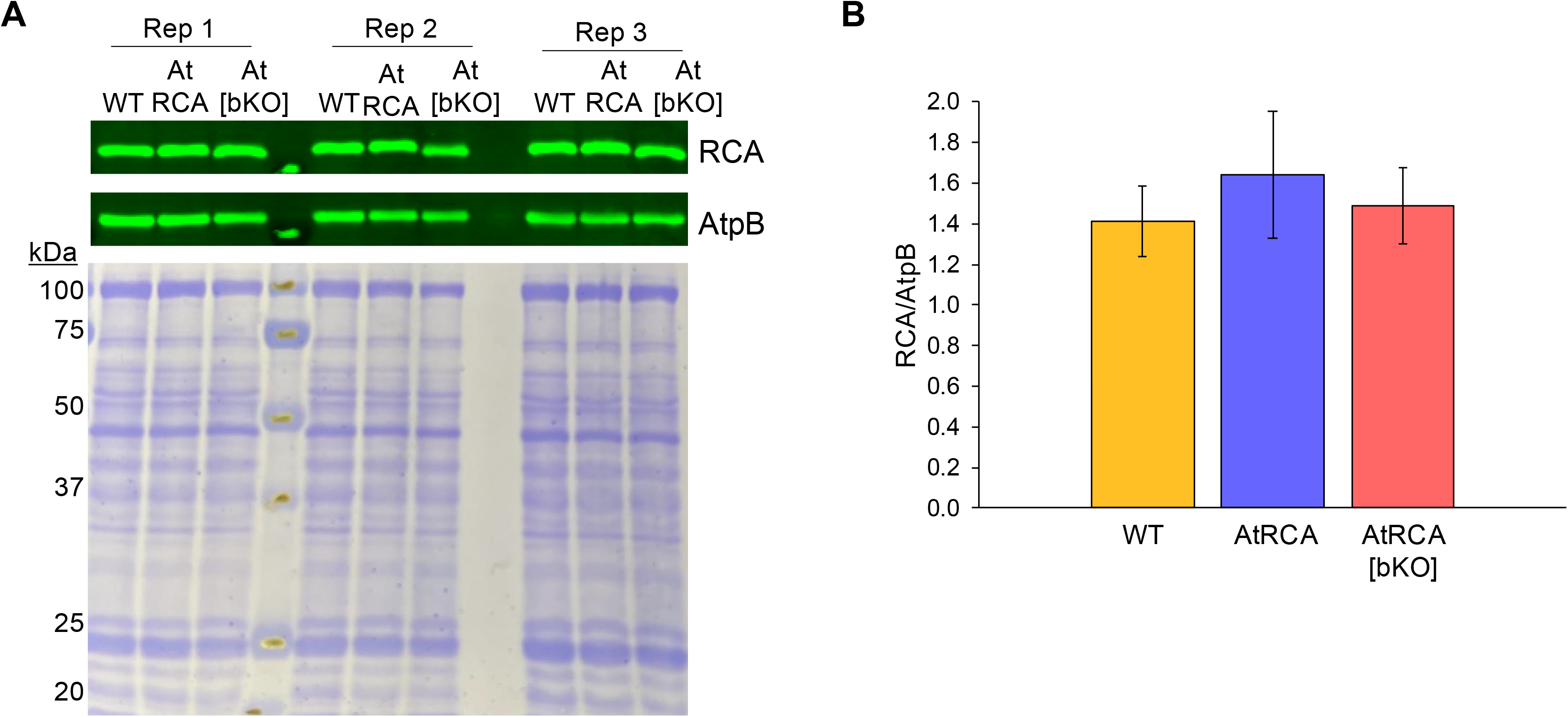
Immunoblot analysis of total protein isolated from WT and transgenic lines (event 1). (A) Protein was extracted from three biological replicates of each genotype, separated by 10% SDS-PAGE and probed with the indicated antibodies (right). Immunoblots were scanned on a Licor Odyssey and a Coomassie brilliant blue (CBB) stained membrane is below to reflect loading with molecular mass indicated on the left (kDa). (B) Quantification of total βRCA relative to AtpB (ATP synthase β subunit; n=3). Error bars are ± SD.

**Fig. S7.**
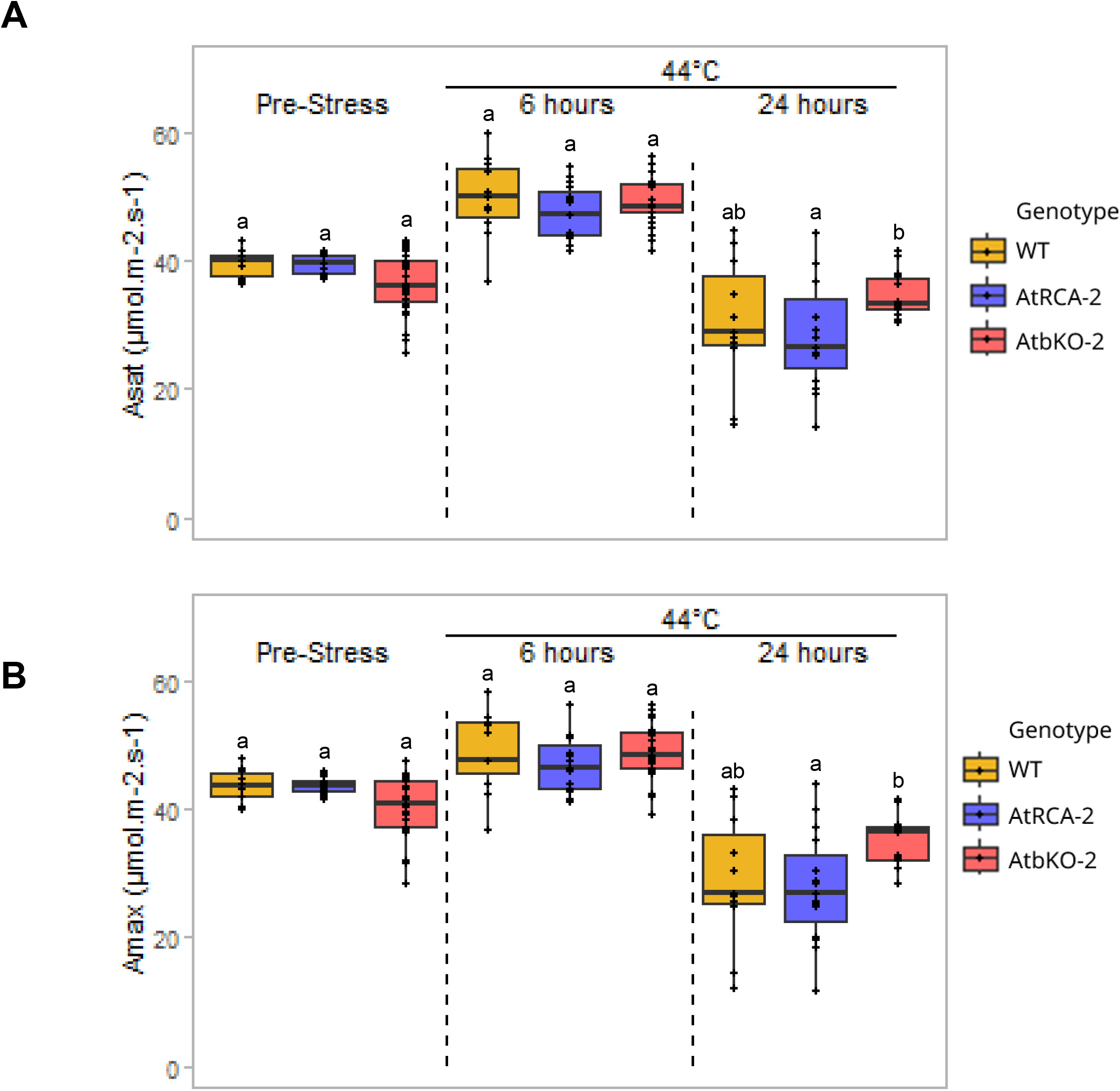
CO_2_ assimilation from a second Agave transgenic event compared to WT. Measurements were made at saturating light (A; 1500 µmol m^-2^ s^-1^ photosynthetic photon flux density (PPFD)) and saturating light and maximum CO_2_ (B). Plants were grown at 25°C for 25-27 days (pre-stress), after which they were put at 44°C and CO_2_ assimilation was measured after 6 h and 24 h (n=6-11 for WT, 14-17 for AtRCA and 16-28 for AtRCA [bKO]). Box plot descriptions are per Fig. 4. and different lowercase letters show significant differences (P<0.05) determined by two-way ANOVA with Tukey’s HSD test.

**Fig. S8.**
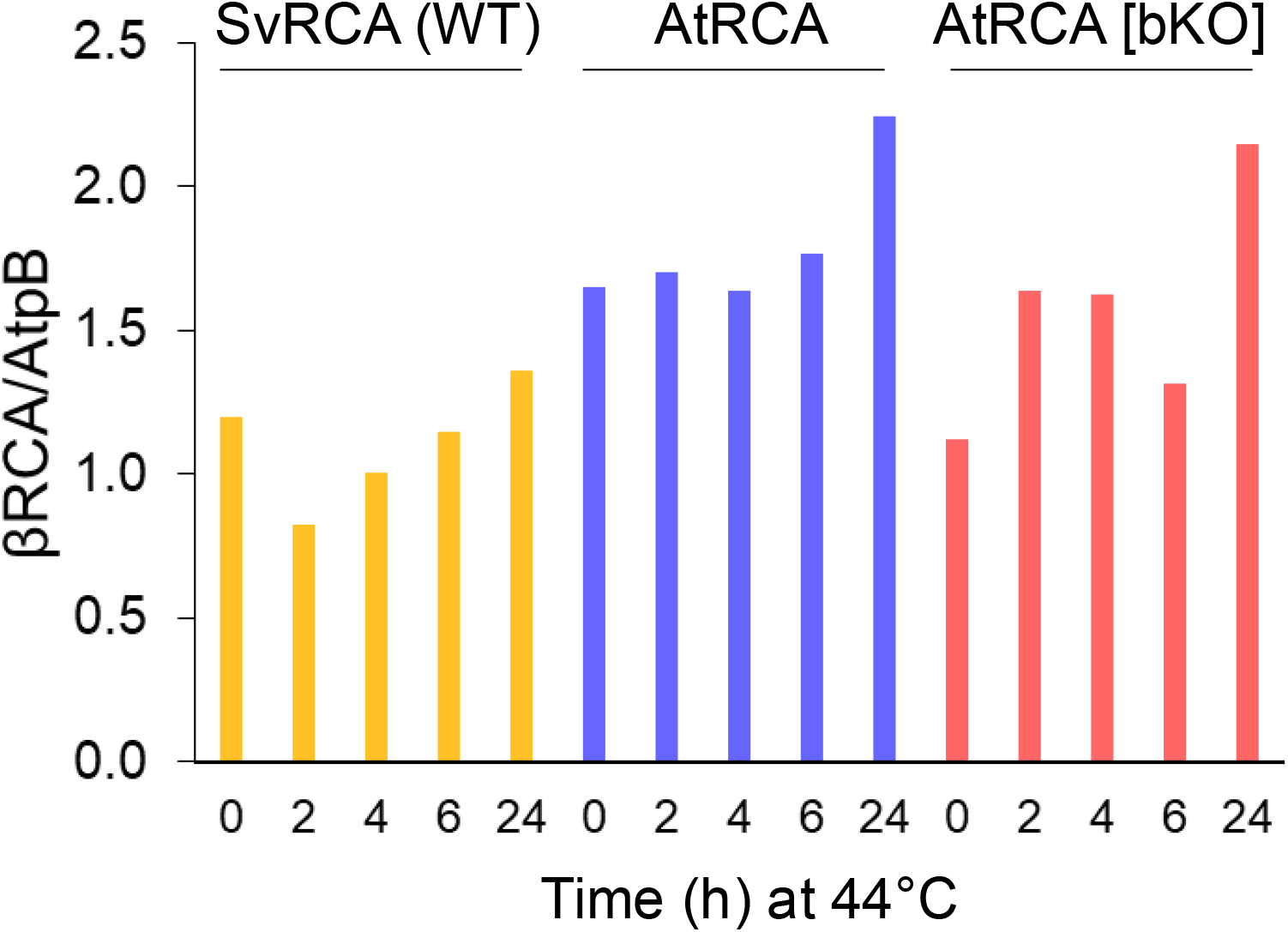
Quantification of the immunoblot analysis in Fig. 6 of RCAβ relative to AtpB using ImageStudio lite. Tissue samples from the youngest fully expanded leaf were collected after plants were incubated at 44°C for the time indicated. Total protein was separated in an 8% SDS-PAGE on an equal area basis for each sample, transferred to PVDF membrane, probed with the relavent antibody, then scanned on the Licor Odyssey.

**Fig. S9.**
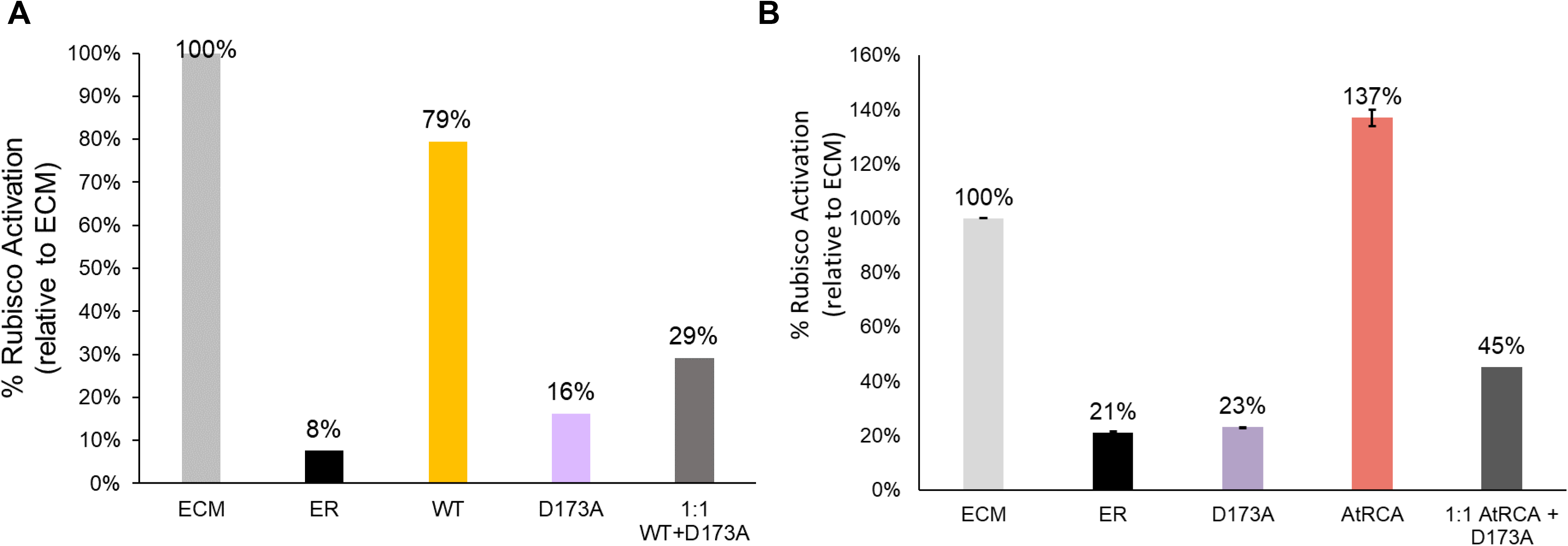
Rubisco activation with and without the Setaria RCA ATPase mutant, D173A, with Asp substituted for Ala at position 173 which inactivates the ATPase catalytic site. (A) Activation of Rubisco by Setaria (WT) RCA, D173A or a 1:1 mass ratio of WT and D173A after 3 min at 25°C using 6.4 mg/mL RCA. (B) Agave (At) RCA, D173A or a 1:1 mass ratio of AtRCA and D173A after 6 minutes at 25°C using 5 mg/mL RCA. Data reflect the ability of the RCA isoform to activate fully inhibited Rubisco (ER) relative to fully activated Rubisco (ECM). Data represent n=1-2 biological replicates with at least 3 technical replicates each with error bars ± SE. D173A was cloned into pHUE, expressed and purified *in vitro* per the manuscript methods.

**Table S1.**
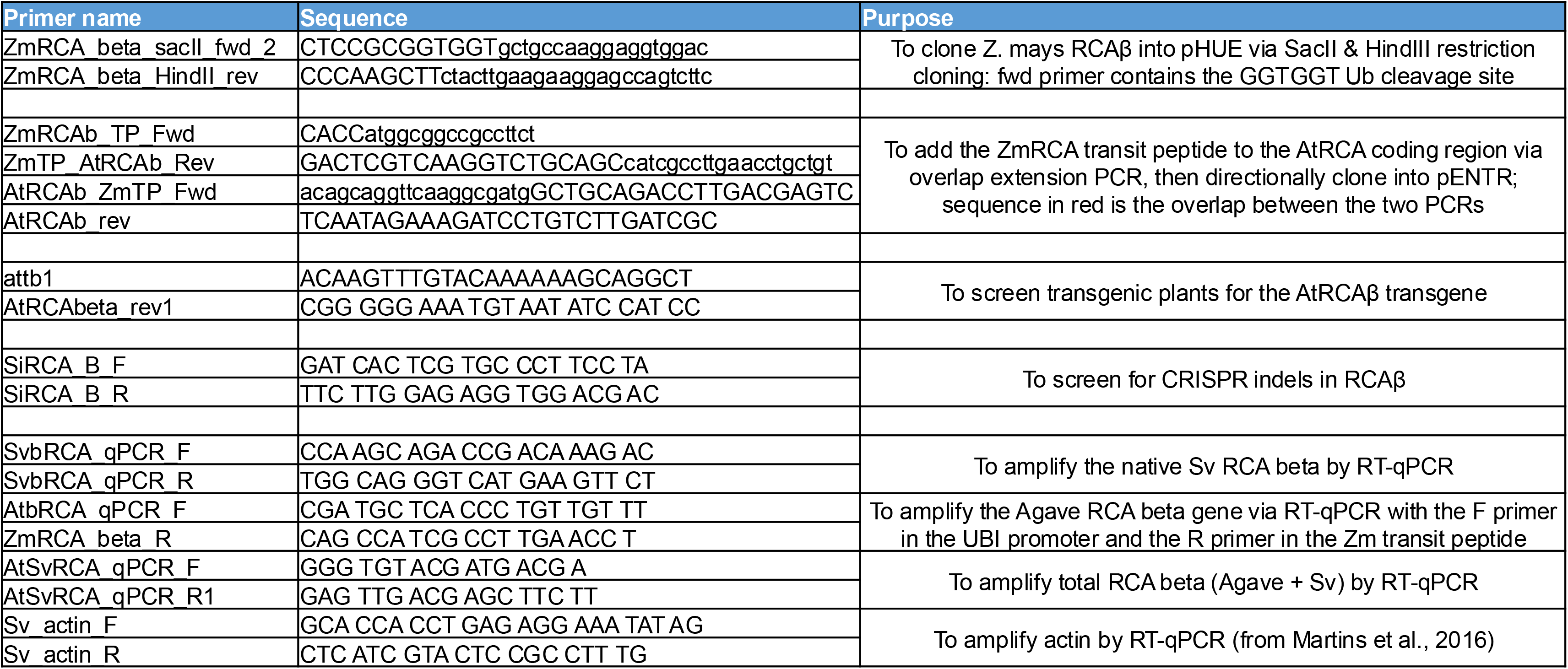
Primers used in this study.

## References

1. Barta C, Dunkle AM, Wachter RM, Salvucci ME. 2010. Structural changes associated with the acute thermal instability of Rubisco activase. Archives of Biochemistry and Biophysics 499, 17– 25.

2. Bhat JY, Thieulin-Pardo G, Hartl FU, Hayer-Hartl M. 2017. Rubisco Activases: AAA+ Chaperones Adapted to Enzyme Repair. Front Mol Biosci 4, 20.

3. Bi H, Liu P, Jiang Z, Ai X. 2017. Overexpression of the rubisco activase gene improves growth and low temperature and weak light tolerance in Cucumis sativus. Physiologia Plantarum 161, 224–234.

4. von Caemmerer S, Hendrickson L, Quinn V, Vella N, Millgate AG, Furbank RT. 2005. Reductions of Rubisco activase by antisense RNA in the C4 plant Flaveria bidentis reduces Rubisco carbamylation and leaf photosynthesis. Plant physiology 137, 747–55.

5. Carmo-Silva E, Salvucci ME. 2013. The Regulatory Properties of Rubisco Activase Differ among Species and Affect Photosynthetic Induction during Light Transitions. Plant Physiology 161, 1645–1655.

6. Catanzariti A-M, Soboleva TA, Jans DA, Board PG, Baker RT. 2004. An efficient system for high-level expression and easy purification of authentic recombinant proteins. Protein Science 13, 1331–1339.

7. Crafts-Brandner SJ, Salvucci ME. 2002. Sensitivity of photosynthesis in a C4 plant, maize, to heat stress. Plant Physiol. 129, 1773–1780.

8. Demirevska-Kepova K, Holzer R, Simova-Stoilova L, Feller U. 2005. Heat stress effects on ribulose-1,5-bisphosphate carboxylase/oxygenase, Rubisco binding protein and Rubisco activase in wheat leaves. Biol. Plant. 49, 521–525.

9. Esquivel MG, Genkov T, Nogueira AS, Salvucci ME, Spreitzer RJ. 2013. Substitutions at the opening of the Rubisco central solvent channel affect holoenzyme stability and CO2/O2 specificity but not activation by Rubisco activase. Photosynth. Res. 118, 209–218.

10. Feng Y, Wu H, Liu H, He Y, Yin Z. 2023. Effects of OsRCA Overexpression on Rubisco Activation State and Photosynthesis in Maize. doi: 10.3390/plants12081614.

11. Friso G, Majeran W, Huang M, Sun Q, van Wijk KJ. 2010. Reconstruction of metabolic pathways, protein expression and homeostasis machineries across maize bundle sheath and mesophyll chloroplasts; large scale quantitative proteomics using the first maize genome assembly. Plant Physiol. 152, 1219–1250.

12. Fukayama H, Ueguchi C, Nishikawa K, Katoh N, Ishikawa C, Masumoto C, Hatanaka T, Misoo S. 2012. Overexpression of rubisco activase decreases the photosynthetic CO2 assimilation rate by reducing rubisco content in rice leaves. Plant Cell Physiol. 53, 976–986.

13. Gjindali A, Page R, Ashton CJ, Robertson I, Page MT, Bloemers D, Gould PD, Worrall D, Orr DJ, Carmo-Silva E. 2025. Two cowpea Rubisco activase isoforms for crop thermotolerance. The New phytologist doi: 10.1111/NPH.70271.

14. Hotto AM, Gartner S, Eshenour K, Stern DB. 2025. The α form of Rubisco Activase supports photosynthesis during heat stress in the absence of the β form in Setaria viridis. Journal of experimental botany doi: 10.1093/JXB/ERAF289.

15. Jiang H, Barbier H, Brutnell T. 2013. Methods for performing crosses in Setaria viridis, a new model system for the grasses. J. Vis. Exp. doi: 10.3791/50527.

16. Kim SY, Slattery RA, Ort DR. 2021. A role for differential Rubisco activase isoform expression in C4 bioenergy grasses at high temperature. GCB Bioenergy 13, 211–223.

17. Kumar A, Li C, Portis Jr. AR. 2009. Arabidopsis thaliana expressing a thermostable chimeric Rubisco activase exhibits enhanced growth and higher rates of photosynthesis at moderately high temperatures. Photosynth Res 100, 143–153.

18. Kurek I, Chang TK, Bertain SM, Madrigal A, Liu L, Lassner MW, Zhu G. 2007. Enhanced thermostability of Arabidopsis Rubisco activase improves photosynthesis and growth rates under moderate heat stress. Plant Cell 19, 3230–3241.

19. Van De Loo FJ, Salvucci ME. 1998. Involvement of two aspartate residues of rubisco activase in coordination of the ATP γ-phosphate and subunit cooperativity. Biochemistry 37, 4621–4625.

20. Mahajan S, Thakur P, Das S, Sharma RP, Manuja S, Jha PK, Saini A, Sahoo C, Fayezizadeh MR. 2025. Impression of contemporary heat stress complexities in agricultural crops: a review. Plant Growth Regulation 2025 105:6 105, 1805–1823.

21. Mann DGJ, Lafayette PR, Abercrombie LL, et al. 2012. Gateway-compatible vectors for high-throughput gene functional analysis in switchgrass (Panicum virgatum L.) and other monocot species. Plant Biotechnology Journal 10, 226–236.

22. Markelz NH, Costich DE, Brutnell TP. 2003. Photomorphogenic responses in maize seedling development. Plant Physiol. 133, 1578–1591.

23. Masumoto C, Fukayama H, Hatanaka T, Uchida N. 2012. Photosynthetic Characteristics of Antisense Transgenic Rice Expressing Reduced Levels of Rubisco Activase. Plant Production Science 15, 174–182.

24. Mate CJ, Von Caemmerer S, Evans JR, Hudson GS, Andrews T. 1996. The relationship between CO2-assimilation rate, Rubisco carbamylation and Rubisco activase content in activase-deficient transgenic tobacco suggests a simple model of activase action. Planta 198, 604–613.

25. Mate CJ, Hudson CS, Von Caemmerer S, Evans JR, Andrews TJ. 1993. Reduction of ribulose bisphosphate carboxylase activase levels in tobacco (Nicotiana tabacum) by antisense RNA reduces ribulose bisphosphate carboxylase. Plant Physiol. 102, 1119–1128.

26. Nagarajan R, Kahlon KS, Mohan A, Gill KS. 2025. Tandemly duplicated Rubisco activase genes of cereals show differential evolution and response to heat stress. Plant Molecular Biology 115, 1–16.

27. Ogbaga CC, Stepien P, Athar HU, Ashraf M. 2018. Engineering Rubisco activase from thermophilic cyanobacteria into high-temperature sensitive plants. Crit Rev Biotechnol 38, 559– 572.

28. Oh ZG, Robison TA, Loh DH, Ang WSL, Ng JZY, Li FW, Gunn LH. 2024. Unique biogenesis and kinetics of hornwort Rubiscos revealed by synthetic biology systems. Molecular Plant 17, 1833–1849.

29. Pak J-H, Seo B, Dal Song S. 1997. Ultrastructural Aspects of Kranz Anatomy in Digitaria sanguinalis and Setaria viridis (Poaceae). J. Plant Biol 40, 102–109.

30. Perdomo JA, Capó-Bauçà S, Carmo-Silva E, Galmés J. 2017. Rubisco and rubisco activase play an important role in the biochemical limitations of photosynthesis in rice, wheat, and maize under high temperature and water deficit. Front. Plant Sci. 8.

31. Prado K, Xue B, Johnson JE, Field S, Stata M, Hawkins CL, Hsia R-C, Liu H, Cheng S, Rhee SY. 2025. Photosynthetic acclimation is a key contributor to exponential growth of a desert plant in Death Valley summer. Current Biology doi: 10.1016/J.CUB.2025.10.004.

32. Qin K, Ye X, Luo S, Fernie AR, Zhang Y. 2025. Engineering carbon assimilation in plants. Journal of Integrative Plant Biology 67, 926–948.

33. Qu Y, Mueller-Cajar O, Yamori W. 2023. Improving plant heat tolerance through modification of Rubisco activase in C3 plants to secure crop yield and food security in a future warming world. Journal of Experimental Botany 74, 591–599.

34. Qu Y, Sakoda K, Fukayama H, Kondo E, Suzuki Y, Makino A, Terashima I, Yamori W. 2021. Overexpression of both Rubisco and Rubisco activase rescues rice photosynthesis and biomass under heat stress. Plant, Cell & Environment 44, 2308–2320.

35. Salvucci ME, Crafts-Brandner SJ. 2004. Relationship between the Heat Tolerance of Photosynthesis and the Thermal Stability of Rubisco Activase in Plants from Contrasting Thermal Environments. Plant Physiology 134, 1460–1470.

36. Salvucci ME, Osteryoung KW, Crafts-Brandner SJ, Vierling E. 2001. Exceptional Sensitivity of Rubisco Activase to Thermal Denaturation in Vitro and in Vivo. Plant Physiology 127, 1053– 1064.

37. Scafaro AP, Atwell BJ, Muylaert S, Reusel B V, Ruiz GA, Rie J V, Galle A. 2018. A thermotolerant variant of Rubisco activase From a wild relative improves growth and seed yield in rice under heat stress. Front Plant Sci 9, 1663.

38. Scafaro AP, Bautsoens N, Boer B Den, Van Rie J, Gallé A. 2019. A Conserved Sequence from Heat-Adapted Species Improves Rubisco Activase Thermostability in Wheat. Plant Physiology 181, 43–54.

39. Sharkey TD, Badger MR, von Caemmerer S, Andrews TJ. 2001. Increased heat sensitivity of photosynthesis in tobacco plants with reduced Rubisco activase. Photosynthesis research 67, 147–56.

40. Shivhare D, Mueller-Cajar O. 2017. In Vitro Characterization of Thermostable CAM Rubisco Activase Reveals a Rubisco Interacting Surface Loop. Plant physiology 174, 1505–1516.

41. Shivhare D, Ng J, Tsai YCC, Mueller-Cajar O. 2019. Probing the rice Rubisco–Rubisco activase interaction via subunit heterooligomerization. Proceedings of the National Academy of Sciences of the United States of America 116, 24041–24048.

42. da Silva GE, Obst S, Carvalho P, Forner J, Ruf S, Saibo NJM, Bock R. 2026. Generation of a recipient line for Rubisco engineering by multiplex genome editing in tobacco. The Plant Journal 126, e70930.

43. Somerville CR, Portis AR, Ogren WL. 1982. A mutant of Arabidopsis thaliana which lacks activation of RuBP Carboxylase in vivo. Plant Physiol. 70, 381–387.

44. Sparrow-Muñoz I, Chen TC, Burgess SJ. 2023. Recent developments in the engineering of Rubisco activase for enhanced crop yield. Biochemical Society Transactions 51, 627–637.

45. Stainbrook SC, Aubuchon LN, Chen A, Johnson E, Si A, Walton L, Ahrendt AJ, Strenkert D, Jez JM. 2024. C4 grasses employ distinct strategies to acclimate rubisco activase to heat stress. Bioscience reports 44.

46. Suganami M, Suzuki Y, Tazoe Y, Yamori W, Makino A. 2021. Co-overproducing Rubisco and Rubisco activase enhances photosynthesis in the optimal temperature range in rice. Plant physiology 185, 108–119.

47. Thielen PM, Pendleton AL, Player RA, Bowden K V., Lawton TJ, Wisecaver JH. 2020. Reference Genome for the Highly Transformable Setaria viridis ME034V. G3 Genes|Genomes|Genetics 10, 3467–3478.

48. Waheeda K, Kitchel H, Wang Q, Chiu PL. 2023. Molecular mechanism of Rubisco activase: Dynamic assembly and Rubisco remodeling. Frontiers in Molecular Biosciences 10.

49. Wang Y, Chan KX, Long SP. 2021. Towards a dynamic photosynthesis model to guide yield improvement in C4 crops. The Plant Journal 107, 343–359.

50. Wang XX, Huang CH, Morales-Briones DF, et al. 2024. Phylotranscriptomics reveals the phylogeny of Asparagales and the evolution of allium flavor biosynthesis. Nature Communications 2024 15:1 15, 9663-.

51. Wang ZY, Snyder GW, Esau BD, Portis AR, Ogren WL. 1992. Species-dependent variation in the interaction of substrate-bound Ribulose-1,5-Bisphosphate Carboxylase/Oxygenase (Rubisco) and Rubisco Activase. Plant Physiol. 100, 1858–1862.

52. Wijewardene I, Mishra N, Sun L, Smith J, Zhu X, Payton P, Shen G, Zhang H. 2020. Improving drought-, salinity-, and heat-tolerance in transgenic plants by co-overexpressing Arabidopsis vacuolar pyrophosphatase gene AVP1 and Larrea Rubisco activase gene RCA. Plant Science 296, 110499.

53. Wijewardene I, Shen G, Zhang H. 2021. Enhancing crop yield by using Rubisco activase to improve photosynthesis under elevated temperatures. Stress Biology 1, 1–20.

54. Wu T, Yu J, Gale-Day Z, Woo A, Suresh A, Hornsby M, Gestwicki JE. 2020. Three Essential Resources to Improve Differential Scanning Fluorimetry (DSF) Experiments. bioRxiv doi: 10.1101/2020.03.22.002543.

55. Yamori W, von Caemmerer S. 2009. Effect of Rubisco Activase Deficiency on the Temperature Response of CO2 Assimilation Rate and Rubisco Activation State: Insights from Transgenic Tobacco with Reduced Amounts of Rubisco Activase. Plant Physiology 151, 2073–2082.

56. Yamori W, Inagaki M, Fukayama H. 2026. Overexpression of Rubisco Activase Improves Photosynthesis and Plant Growth in Rice Under Fluctuating Light Under Future High CO2 Conditions. Plant, Cell & Environment doi: 10.1111/PCE.70617.

57. Yang M, Wang X, Zhang X, Wei X, Guo J. 2026. Overexpression of the halophyte Suaeda salsa Rubisco activase gene SsRCA in Arabidopsis improves plant photosynthesis under salt-stressed conditions. Journal of Plant Physiology 316, 154670.

58. Zhang N, Schürmann P, Portis AR. 2001. Characterization of the regulatory function of the 46-kDa isoform of Rubisco activase from Arabidopsis. Photosynthesis Research 68, 29–37.

59. Zhang P, Sharwood RE, Carroll A, Estavillo GM, von Caemmerer S, Furbank RT. 2025. Systems analysis of long-term heat stress responses in the C4 grass Setaria viridis. The Plant Cell 37, 5.

